# Dynamics of microRNA expression during mouse prenatal development

**DOI:** 10.1101/492918

**Authors:** Sorena Rahmanian, Rabi Murad, Alessandra Breschi, Weihua Zeng, Mark Mackiewicz, Brian Williams, Carrie Davis, Brian Roberts, Sarah Meadows, Dianna Moore, Diane Trout, Chris Zaleski, Alex Dobin, Lei-Hoon Sei, Jorg Drenkow, Alex Scavelli, Thomas Gingeras, Barbara Wold, Richard M. Myers, Roderic Guigó, Ali Mortazavi

**Affiliations:** Department of Developmental and Cell Biology, University of California Irvine, Irvine, CA 92697, USA; Center for Complex Biological Systems, University of California Irvine, Irvine, CA 92697, USA; Bioinformatics and Genomics, Centre for Genomic Regulation (CRG) and UPF, Barcelona 08003, Catalonia, Spain; Division of Biology, California Institute of Technology, Pasadena, California 91125, USA; HudsonAlpha Institute for Biotechnology, Huntsville, AL 35806, USA; Functional Genomics, Cold Spring Harbor Laboratory, Cold Spring Harbor, New York 11724, USA

## Abstract

MicroRNAs (miRNAs) play a critical role as post-transcriptional regulators of gene expression. The ENCODE project profiled the expression of miRNAs in a comprehensive set of tissues during a time-course of mouse embryonic development and captured the expression dynamics of 785 miRNAs. We found distinct tissue and developmental stage specific miRNA expression clusters, with an overall pattern of increasing tissue specific expression as development proceeds. Comparative analysis of conserved miRNAs in mouse and human revealed stronger clustering of expression patterns by tissue types rather than by species. An analysis of messenger RNA gene expression clusters compared with miRNA expression clusters identifies the potential role of specific miRNA expression clusters in suppressing the expression of mRNAs specific to other developmental programs in the tissue where these microRNAs are expressed during embryonic development. Our results provide the most comprehensive timecourse of miRNA expression as an integrated part of the ENCODE reference dataset for mouse embryonic development.

## INTRODUCTION

Development is a well-orchestrated process primarily controlled by transcriptional regulators with post-transcriptional regulators such as microRNAs (miRNAs) playing an essential role in fine tuning gene expression dynamics. MicroRNAs are small ~22 nucleotide (nt) endogenous non-coding RNAs that regulate gene expression by mediating the post-transcriptional degradation of messenger RNA (mRNA) or by hindering the translation of proteins (Bartel, 2004; He & Hannon, 2004). Their biogenesis occurs in several steps, starting with transcription of typically polyadenylated primary miRNA (pri-miRNA) transcripts (>200 nt), sometimes referred to as the “host genes”. These pri-miRNA have a characteristic hairpin structure that is cleaved in the nucleus by the enzyme Drosha into pre-miRNA (~60 nt), which are exported to the cytoplasm before finally being processed into 21-24 nt mature miRNA by the enzyme Dicer (Han et al., 2006). The first miRNA was discovered in the nematode *C. elegans* as perturbing its cell developmental lineage (Lee, Feinbaum, & Ambros, 1993) and since then thousands of miRNAs have been discovered in diverse plants, metazoans, and some viruses (Kozomara & Griffiths-Jones, 2011).

Many studies have shown that the deletion of key players in the biogenesis of miRNA such as Ago2, Dicer1 and Dgcr8 will lead to embryonic lethality and arrest (Alisch, Jin, Epstein, Caspary, & Warren, 2007; Bernstein et al., 2003; Morita et al., 2007; Wang, Medvid, Melton, Jaenisch, & Blelloch, 2007). However loss of single miRNAs does not have as dramatic an effect as knocking out all the miRNAs in the organism (Park, Choi, & McManus, 2010). This could be due to the redundancy of miRNA-mRNA interactions as each mRNA could be targeted by multiple miRNAs and thus the lack of one miRNA would be compensated by others. Hence there is a strong rationale for studying the role of miRNAs as a functional group or unit. Studies have shown that most genes are potential targets of miRNAs (Friedman, Farh, Burge, & Bartel, 2009) and that miRNAs are involved in regulating diverse cellular processes during development and homeostasis (Vidigal & Ventura, 2015). Dysregulation of miRNA expression is known to underlie numerous diseases and developmental defects such as cancer (Lin & Gregory, 2015), cardiovascular diseases (Romaine, Tomaszewski, Condorelli, & Samani, 2015; Zhao et al., 2015), and neurological diseases (Cao, Li, & Chan, 2016).

MicroRNAs have been profiled in various tissues and primary cells in diverse metazoans and plants (Ehrenreich & Purugganan, 2008; Lagos-Quintana et al., 2002; Wienholds et al., 2005). Mineno and colleagues used massively parallel signature sequencing (MPSS) technology to profile miRNAs in mouse whole embryos during three embryonic stages (e9.5, e10.5, and e11.5) and were able to detect 390 distinct miRNAs (Mineno et al., 2006). Chiang and colleagues extended this work by sequencing small RNAs from mouse brain, ovary, testes, embryonic stem cells, embryonic stages of complete embryos from three developmental stages, and whole newborns to profile the expression of 398 annotated and 108 novel miRNAs (Chiang et al., 2010). Landgraf and colleagues cloned and sequenced more than 250 small RNA libraries from 26 different organs and cell types from humans and rodents to profile miRNA expression and describe various other miRNA characteristics (Landgraf et al., 2007). More recently, the FANTOM5 project has created a miRNA expression atlas using deep-sequencing data from 396 human and 47 mouse RNA samples (De Rie et al., 2017); however, many of these mouse samples were simply replicates of a handful of mouse cells lines with and without stimulation. Previous efforts by the ENCODE Consortium affiliates focused on a meta-analysis of previously published 501 human and 236 mouse small RNA sequencing data sets from a multitude of sources to characterize splicing-derived miRNAs (mirtrons) in the human and mouse genomes (Ladewig, Okamura, Flynt, Westholm, & Lai, 2012). However, the diversity of the source tissues and the different underlying experimental protocols from the disparate primary sources complicated any sort of systematic quantitative analysis. Last but not least, many individual studies have focused on the expression of particular microRNAs in certain tissues in a handful of (typically 2-3) mouse developmental timepoints. Therefore, a complete and systematic atlas of miRNA expression during development in tissues representative of the major organ systems and broad number of mouse embryonic stages is still missing. This is not only helpful for understanding mouse development, but also for studying the potential role of microRNAs in human development where access to the same timepoints is either very difficult or outright impossible.

With the growing evidence of the critical role of miRNAs in homeostasis and disease, multiple techniques have been developed for profiling the expression of mature miRNAs, each with their own strengths (Mestdagh et al., 2014). RNA-seq typically refers to the profiling of expressed transcripts 200 nt or longer including the messenger RNAs (mRNA) and long non-coding RNAs (lncRNA) (Mortazavi, Williams, McCue, Schaeffer, & Wold, 2008), which in this work we will refer to as messenger RNA-seq (mRNA-seq), whereas there are also multiple miRNA-specific sequencing protocols such as microRNA-seq (Roberts et al., 2015) and short RNA-seq (Fejes-Toth et al., 2009). There are also hybridization-based assays such as microarrays as well as molecule counting such as NanoString, which involves hybridization of color-coded molecular barcodes (Geiss et al., 2008; Wyman et al., 2011). As mature miRNAs are processed from longer host pri-miRNAs and the annotated pri-miRNAs are predominantly protein-coding or lncRNA transcripts (Cai, Hagedorn, & Cullen, 2004), we hypothesize that mRNA-seq should be able to profile the expression of pri-miRNAs. However, there is a significant number of miRNAs whose host genes have not been characterized yet. Furthermore, an important question is whether the expression of pri-miRNAs can reliably predict the expression of their corresponding mature miRNAs. This would allow the simultaneous profiling of mature miRNA expression along with mRNAs using mRNA-seq. Previous studies have attempted to answer this question in specific cell types (Zeng et al., 2016). Availability of matching mRNA-seq and microRNA-seq data sets for the same samples in our study provides a unique opportunity to answer this question. Furthermore, the corresponding mRNA data can shed light into the targeting of these miRNAs and their functional role during the embryonic development.

Each miRNA targets a set of mRNAs through Watson-Crick pairing between miRNA seed region (positions 2-7 from 5’ end) and the binding sites on their targets (Bartel, 2009). Such complementary base-pairing has been used to computationally predict miRNA targets (Bartel, 2009). Expression of miRNAs and mRNAs in matching samples have been used to identify miRNA-mRNA interactions, for example in cancer (McLendon, 2008). Several methods such as biclustering (Jin & Lee, 2015) have been used to infer miRNA-mRNA interactions from gene expression data. However, the expression levels of mRNAs are often affected by multiple factors and comparison of mRNA and miRNA expressions cannot establish a functional relationship by itself. Therefore, an approach that integrates miRNA and mRNA expression data and their predicted interactions should provide better inference of their functional interaction networks.

In this study, we have used microRNA-seq and NanoString to characterize the expression patterns of known microRNAs using a set of 16 different mouse tissues at 8 embryonic (e10.5 – P0) stages that were specifically selected by the ENCODE consortium for a wide variety of functional sequencing assays such as mRNA-seq, ChIP-seq and DNase-seq. The value of this dataset is that the tissue samples and stages are all matched. We show one example of integrative analysis of the microRNA-seq data with matching ENCODE mRNA-seq (and ChIP-seq) data to compare the characteristics and dynamics of miRNA expression to characterize the changes in overall tissue specificity of particular microRNAs during mouse development. In particular, we compute enrichment of computationally predicted miRNA targets in certain mouse tissues along with the negative partial correlation analysis of miRNA and mRNA expression clusters during mouse development to identify developmental processes targeted by miRNAs and show that groups of miRNAs expressed in one or more tissue target groups of developmentally important mRNAs highly expressed in other tissues.

## RESULTS

### A reference miRNA catalog across mouse development

As part of the ENCODE project, we used microRNA-seq and NanoString to profile mature miRNAs during mouse embryonic development and matched it to mRNA-seq in order to profile the expression of pri-miRNAs (Supplementary Fig 1a). This study encompasses 156 microRNA-seq and 154 NanoString datasets in matching mouse tissues with two biological replicates each (Fig 1a). We found a high correlation between microRNA-seq and NanoString data in the same tissue at the same time point (median Spearman correlation = 0.68) (Supplementary Fig. 1b), which matches the reproducibility between platforms reported previously (Mestdagh et al., 2014). All data from this study are available from the ENCODE data portal (www.encodeproject.org) with the accession numbers listed in Supplementary Table 1.

**Figure 1:**
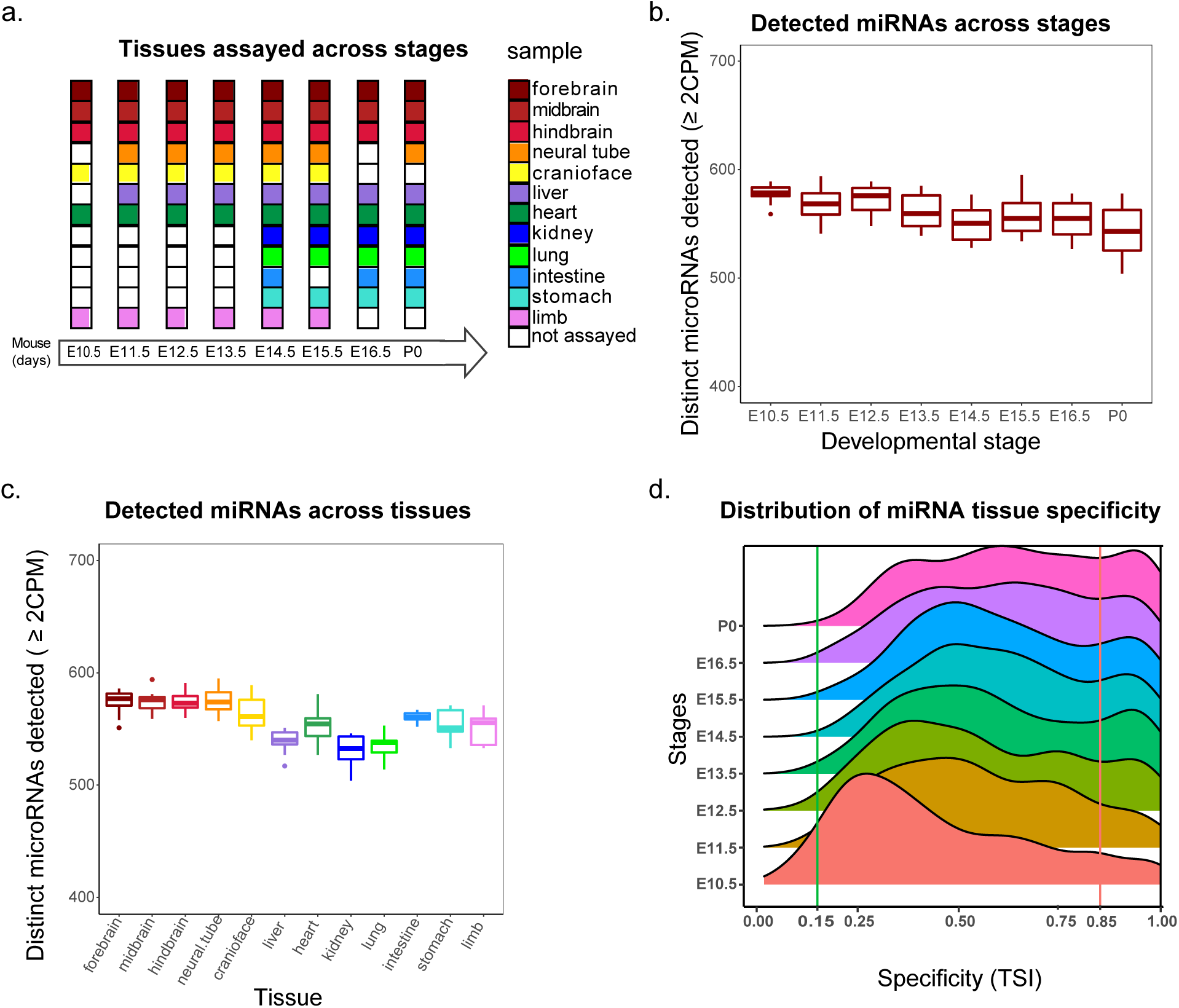
Overview of mouse ENCODE miRNA data sets. **(a)** Primary tissues representative of major organ systems were profiled in a time course of mouse embryonic development, **(b)** Number of distinct miRNAs detected in different developmental stages (minimum 2 CPM). **(c)** Number of distinct miRNAs detected in tissues (minimum 2 CPM). **(d)** The distribution of tissue specificity of miRNAs expressed at each developmental stage measured as tissue specific index (TSI). The miRNAs are significantly more tissue specific at stage of P0 compared to E10.5 (p-value < 2.2e-16).

We used a set of three spike-ins of different sequence lengths (22 bp, 25 bp, and 30 bp) in decreasing concentrations (5000, 500, and 50 pM respectively) in our microRNA-seq samples to assess replicate concordance for different library normalization strategies (Supplementary Fig 1c). While the spike-in counts were highly concordant for biological replicates for each sample, they differ for different stages of mouse embryonic development using counts per million (CPM) normalization only. We found that TMM normalization of miRNA CPMs ameliorates such differences in spike-in expression across developmental stages. Therefore, we normalized our data using TMM normalization for downstream analysis.

We used microRNA-seq reads to quantify miRNA expression levels using miRBase version 22 annotations, which includes 1981 mature miRNAs. We detected 785 of these mature miRNAs expressed in at least one of the samples; About 80% of these mature miRNAs correspond to the pre-miRNAs identified as highly confident by miRBase criteria (Kozomara et al., 2013). This cohort of miRNAs encompasses 61% of highly-confidence miRNAs in miRBase compared to the 65% recovered by FANTOM dataset (Derek et al., 2017). There are no significant differences in the number of distinct miRNAs expressed in mouse tissues and developmental stages and although stage P0 has the highest number of tissues profiled as well as the least number of distinct miRNAs detected (Fig 1b). This result is in contrast to the finding that the absolute numbers of expressed miRNAs increase over the developmental time in other model organisms such as *Drosophila melanogaster* (Ninova, Ronshaugen, Griffiths-jones, & Griffiths-jones, 2014). At the tissue level, we find that the nervous system samples show the highest number of distinct miRNAs expressed (Fig 1c).

### Dynamics of miRNA tissue specificity during development

As previous studies have shown, a few highly expressed miRNAs are responsible for most of the detected expression (Lagos-Quintana et al., 2002; Landgraf et al., 2007), with about 50% of the expression corresponding to the top 10 expressed miRNAs (Supplementary Fig 2). Only 42 miRNAs fall within the top ten expressed miRNAs across our 72 distinct tissue-stage samples. Six of these miRNAs are in the top 10 expressed list for more than half of the samples with miR-16-5p and miR-26a-5p being one of the top expressed miRNAs in every single experiment. To study the specificity of the miRNAs at each stage, we used the Tissue Specificity Index (TSI) as defined previously (Ludwig et al., 2016); using this metric, we found that 40% of the top expressed miRNAs are tissue specific in at least one of the stages that they are highly expressed in. These miRNAs include: miR-1a-3p, miR-208b-3p and miR-351-5p in heart (the last one is only specific in the earlier stages); miR-9-3p, miR-9-5p, miR-124-3p, miR125b-5p and miR92b-5p in brain; miR-122-5p and miR-142a-3p in liver; miR-10a-5p in kidney; miR-194-5p in intestine; miR-196b-5p in limb. (Supplementary Fig 3)

While there are few miRNAs that are expressed ubiquitously (TSI < 0.15) at the earlier stages of embryonic development, most miRNAs become more tissue-specific as the embryo develops further and the landscape of miRNAs shifts from ubiquitousness to being specific (Fig 1d). This shift is partly due to changes in the specificity of the miRNAs throughout development with the following miRNAs showing the most change: miR-128-3p, miR-181a-1-3p, miR-138-5p and miR-3099-3p in brain; miR-101a-3p and miR-496a-3p in liver; and miR-140-5p in kidney (Supplementary Fig 4); all of these miRNAs increase in their specificity from being almost ubiquitous to become tissue specific. However, there is a group of more than 20 miRNAs that stay mostly specific throughout the developmental time points captured in our studies. This group includes some of the well-studied tissue-specific miRNAs such as: miR-9 and miR-92 in brain; miR-1a-3p, miR-208a-3p and miR-133a-3p in heart; miR-122-5p in liver (Supplementary Fig 5). Finally, there is a group of miRNAs that are present in almost all the tissues at every stage of development including: miR-17-3p, miR-421-3p, miR-361-5p and miR-744-5p. (Supplementary Fig 6). In summary, our high-resolution time course captures the distinct patterns of microRNA expression during mouse embryonic development.

### Clustering of microRNAs recovers distinct tissue specific clusters

Global analysis of mouse tissues and developmental stages shows distinct miRNA expression patterns as revealed by principal component analysis (PCA) (Fig 2a). Principal component 1 (PC1) accounts for 24 percent of the variation and clearly separates the various tissues with the nervous system and liver tissues at the extremes, whereas PC2 (15% variation) represents the time component of mouse development with a temporal gradient between early development at embryonic day 10 (e10.5) and postnatal samples right after birth (P0) (Fig 2a), PC3 (10.8% variation) separates kidney samples from liver, PC4 (6.1% variation) separates heart samples from other tissues and PC5 (4.5% variation) separates kidney samples from limb and craniofacial samples. Overall the first five principal components explained over 60% of the variation in the dataset with most of that variation corresponding to specific tissues.

We used maSigPro (Conesa et al., 2013) to cluster the 785 expressed miRNAs based on the tissue-specific changes in their expression during the development. maSigPro identified 535 of these miRNAs as being differentially expressed (Supplementary Table 3) during embryonic development into 16 clusters based on regression of their expression levels in each of the tissues (Fig 2b and Supplementary Fig 7). Clustering of the matching NanoString microRNA data recovers 14 clusters, the majority of which match the microRNA-seq clusters (Fig 2c and Supplementary Fig 8). We focus on the microRNA-seq clusters going forward. Cluster 11 has the highest number of miRNAs in it (96 miRNAs) and these miRNAs are highly expressed in brain. Additionally, the expression of these miRNAs goes up during embryonic development whereas in cluster 2, another brain-specific cluster, the expression of miRNAs goes up initially and comes down after day 14. miRNA clusters 4, 12 and 14 are the second largest clusters, with 54 miRNAs each. Clusters 4 and 12 are composed of miRNAs mostly expressed in liver and heart respectively, whereas miRNAs in cluster 14 are highly expressed in all the tissues except in liver and brain. Analysis of tissue specific miRNAs in each cluster reveals that more than half of the miRNA clusters are enriched for specificity to only one tissue with the rest of clustered enriched for specificity in 2 or 3 different tissues (Supplementary Fig 9). Thus, microRNAs during development show distinct clustered expression in select tissues.

**Figure 2:**
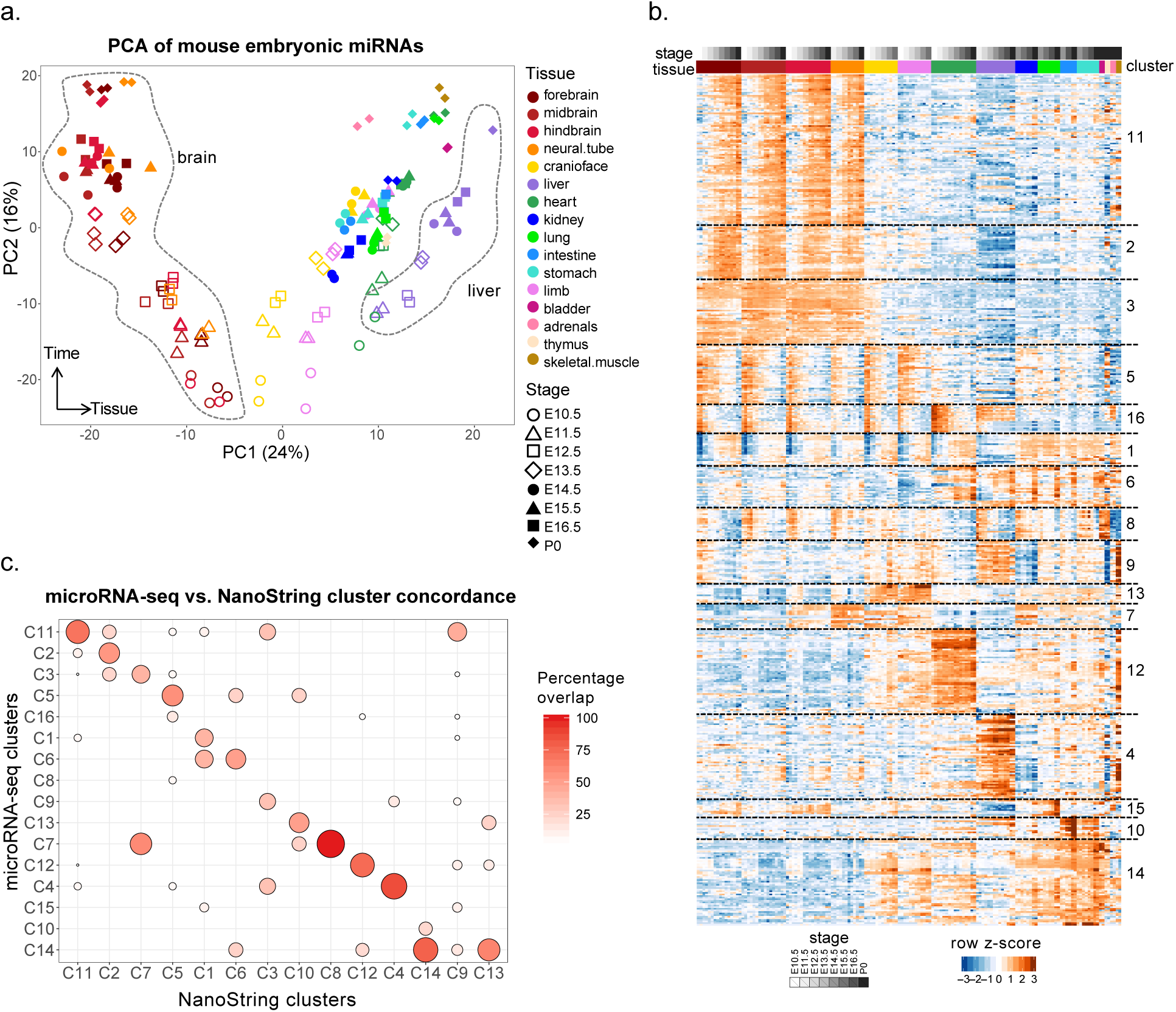
Global properties of mouse miRNA embryonic development time-course. **(a)** Principal component analysis (PCA) of 16 mouse tissues across 8 developmental stages. Tissues are represented by specific colors while shapes denote the various developmental stages, (b) Clustering of miRNAs using maSigPro into 16 non-redundant groups based on median expression profiles (Supplementary Fig 2). (c) Overlap of miRNAs among the 14 clusters of differentially expressed miRNAs (rows) assayed using NanoString (Supplementary figure 3) and the 16 clusters of differentially expressed miRNAs (column) assayed with microRNA-seq. Boxes with increasing intensity of orange color indicate the clusters with increasing orthogonal overlap.

### Comparative dynamics of conserved miRNAs during development

The ENCODE Consortium also collected a limited set of human developmental samples that were characterized with short RNA-seq to profile pre-miRNAs and mature miRNAs for a total of 32 datasets in various tissues during weeks 19-40 of human fetal development that correspond primarily to postnatal day 0 of mouse development (Supplementary Fig 10a). We compared the consistency of microRNA-seq and short RNA-seq in human K562 and GM12878 cells and compared them to datasets from a previous phase of ENCODE. We show that there is high correlation between microRNA-seq and short RNA-seq (Pearson correlation = 0.99) (Supplementary Fig 10b). This level of reproducibility clearly allows us to differentiate between different cell types, even across different methods and batches (Supplementary Fig 10c) and allows us to compare human and mouse microRNA expression levels across the two sequencing techniques.

We quantified human known and novel miRNAs using short RNA-seq and GENCODE v.25 annotations consisting of 1569 known miRNAs supplemented with the novel miRNAs. A global PCA analysis of miRNA expression shows the brain samples clustering as previously seen in mouse (Supplementary Fig 11a). While the availability of human samples was more limited compared to mouse, we identified 279 tissue-specific miRNAs (Supplementary table 4), most of which (83%) are preferentially expressed in neuronal and muscular tissues (Supplementary Fig 11b).

In order to compare miRNA expression across development, we first searched for orthologous miRNAs between mouse and human (Supplementary Fig 12a). We found that a subset of miRNAs is conserved between mouse and human, with 304 miRNAs having a one-to-one orthologous relationship (Supplementary Table 5). Our analysis also revealed that 838 and 516 miRNAs in human and mouse respectively lack a clear ortholog in the other species (Supplementary Fig 12a). We used the set of one-to-one miRNA orthologs to perform PCA on matching tissues in mouse and human, which revealed clustering of samples based on tissue types (Supplementary Fig 12b). Furthermore, comparative clustering of tissues in mouse and human reveals distinct miRNA expression patterns similar to the clustering of tissues in mouse only. As in the mouse developmental time course, the nervous system and liver tissues cluster separately from the rest of the tissues. We compared the sets of tissue-specific orthologous miRNAs across all the available tissues in mouse and human, and represented each comparison as pie charts, where the sizes of the pie charts are in proportion to the number of tissue-specific miRNAs (Supplementary Fig 12c). We found that muscle tissues in mouse and human show the highest conservation (>50%) of expression and that the conservation of expression among the corresponding brain, neural tube and lung tissues is significant (~50%), whereas the conservation of expression between the liver samples is low (Supplementary Fig 12c, Supplementary Table 6). Therefore the conservation of tissue-expression of homologous microRNAs between human and mouse is dependent on the specific tissue.

### Correlation of expression among pri-miRNAs and their corresponding mature miRNAs

The availability of matching microRNA-seq and mRNA-seq data allowed us to evaluate whether the expression of pri-miRNAs is predictive of the expression of their corresponding mature miRNAs. Less than 50% of miRNAs in mouse have annotated primary transcripts in GENCODE version M10 (Supplementary Fig 13). We used mRNA-seq data to assemble additional transcript models and supplement the GENCODE annotations, which increased the number of pri-miRNAs in mouse and human by 7% and 17% percent respectively. A representative novel model transcript in mouse, assembled using all mouse mRNA-seq datasets, overlaps miR-let7a and miR-let7f that were lacking annotated pri-miRNAs and is supported by stage-matched chromatin immunoprecipitation followed by sequencing (ChIP-seq) for both H3K4me3 marking the putative promoter (Supplementary Fig 14) and H3K36me3, as correlate of transcription (Supplementary Fig 13a). Global correlation analysis of the expression levels of the pri-miRNAs (measured by mRNA-seq) and their corresponding miRNAs (measured by microRNA-seq) shows that 143 (41% of the miRNAs expressed at a minimum of 10 CPM) are well correlated (Spearman correlation ≥ 0.6) with their corresponding pri-miRNAs across the developmental time-course. The median Spearman correlation for all the miRNAs and their corresponding pri-miRNAs is 0.51 (Supplementary Fig 13c,d, Supplementary Table 7). Thus, miRNA expression can be imputed from the expression of its primary transcript as measured by mRNA-seq and confirmed using the matching ENCODE ChIP-seq data resources.

### Integrative analysis of microRNA and mRNA expression profiles during mouse development identifies significant anti-correlations of developmentally important genes with microRNAs predicted to target them

In order to understand the connection between microRNAs and the expression of their targets, we developed an integrative analysis pipeline to connect microRNAs to their mRNA targets (Fig 3a). As a first step, we quantified the tissue specificity of miRNA clusters by computing a tissue specificity matrix. The tissue specificity of each miRNA cluster was determined based on the expression changes of miRNAs in each tissue during development. The tissues that had the highest standard deviation of a given miRNA cluster’s expression in different stages were identified as the tissue specificity of that cluster. The tissue-specificity of the miRNA clusters calculated in this manner are highly concordant with the tissue-specificities obtained by the specificity analysis of the individual miRNAs in each cluster. There is at least one miRNA cluster identified for each tissue and at least one tissue identified as tissue specific for each miRNA cluster (Fig 3b, Supplementary Fig 9, Supplementary Table 8).

**Figure 3:**
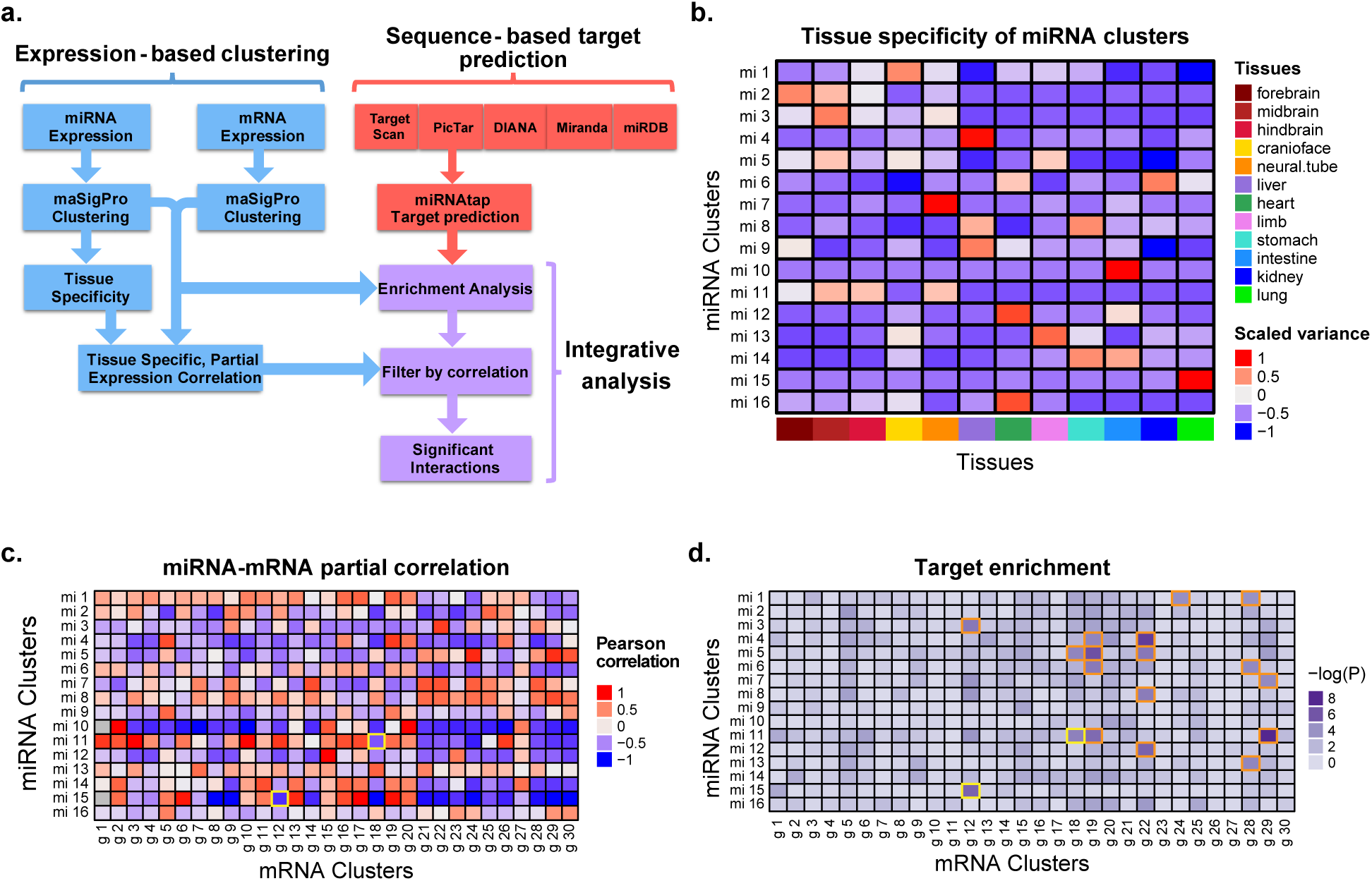
Identification of miRNA-mRNA cluster interactions. **(a)** The potential targets of each miRNA cluster were obtained by applying an ensemble method (miRNAtap) using five different sources and interactions were called significant if they had a negative tissue-specific partial correlation and were enriched beyond the Bonferroni-corrected P-value of 10^−4^. **(b)** Heatmap of the variance of miRNA cluster mean expression during the time course in each tissue. These values were scaled for each cluster separately to identify the tissue specificity of that cluster, **(c)** Tissue-wise partial Pearson correlation between miRNAs clusters and each mRNA clusters identifies significant anti-correlations, **(d)** Heatmap of miRNA cluster target enrichment calculated using χ^2^ statistics. The 18 interactions identified as enriched are boxed in orange and gold. The two gold interactions are further analyzed in Figure 4.

We clustered mouse developmental mRNAs from ENCODE using maSigPro into 30 clusters incorporating 14,827 differentially expressed genes out of the 20,686 genes that were expressed at least once during the development with a replicate average expression of at least two TPM (Supplementary Table 9). About one third of these clusters are specific to a single tissue, with the rest being expressed in multiple tissues. The largest three clusters are clusters 9, 3 and 12 with 1,773, 1,214 and 884 genes in them respectively and all three of these clusters correspond to genes expressed in brain. Most of the tissue specific clusters correspond to liver, heart and lung after the brain.

After identifying the clusters of miRNA and mRNA that are dynamically expressed during development, we calculated the partial correlation between each of the miRNA clusters and each of the mRNA clusters. The partial correlation matrix was built using Pearson correlation between each pair of clusters (miRNA-mRNA) within the context of tissues that the miRNA cluster was active in (using only the miRNA tissue specificity as the context) (Fig 3c, Supplementary Table 10). Using this partial correlation approach, 60% of the miRNA-mRNA cluster interactions are anti-correlated with a mean correlation coefficient value of −0.47. This anti-correlation was used to filter out the positive interactions after target enrichment analysis.

We collected the predicted targets of each of the miRNAs from five different resources and prediction algorithms using miRNAtap (Pajak et al, 2014). We used the unique set of all the predicted targets for miRNAs in each of the miRNA clusters to build a contingency table that contains the distribution of each of these unique target sets among the mRNA clusters. We then performed a χ-square test on the contingency table to study the enrichment of targets in different mRNA clusters (Fig 3d, Supplementary Table 11) and applied a p-value cut off of 0.0001 (Bonferroni corrected P-value: 0.05/ 16*30) to determine the mRNA clusters that were significantly enriched for miRNA cluster targets. 18 interactions between 11 unique miRNA clusters and 7 unique mRNA cluster were identified as significant, however only 9 of these interactions passed the filter for negative partial correlation (Supplementary Fig 15, Supplementary Table 12). The interaction between miRNA cluster 11 and mRNA cluster 18 (Fig 4a, b, and c, Supplementary Table 13) had a P-value of 10^-5^ for the target enrichment and a correlation coefficient of −0.73. The miRNAs involved in this interaction are highly expressed and increase during time in brain whereas the target genes are expressed more highly in other tissues such as limb, cranioface, and heart at the same stages of development. Gene ontology of the targets revealed that this miRNA cluster targets genes involved in the development of skeletal system, cardiac development and vasculature development (p-values < 10^-15^), by presumably downregulating them in the brain. Another interaction between miRNA cluster 15 and mRNA cluster 12 has a P-value of 10^-6^ and negative correlation coefficient of −0.89. This miRNA cluster increases expression mainly in the lung (Fig 4d) whereas the mRNA targets are highly expressed in brain tissues and their expression is very limited in lung (Fig 4e). Gene ontology analysis of this interaction enriches for terms involved with synaptic processes and neuronal systems (p-values of <10^-7^, Fig 4f, Supplementary Table 14). In both of these cases as well as several of the others, the miRNA cluster is enriched for targets that are developmentally important genes for tissues other than the tissue in which the miRNAs are highly expressed.

**Figure 4:**
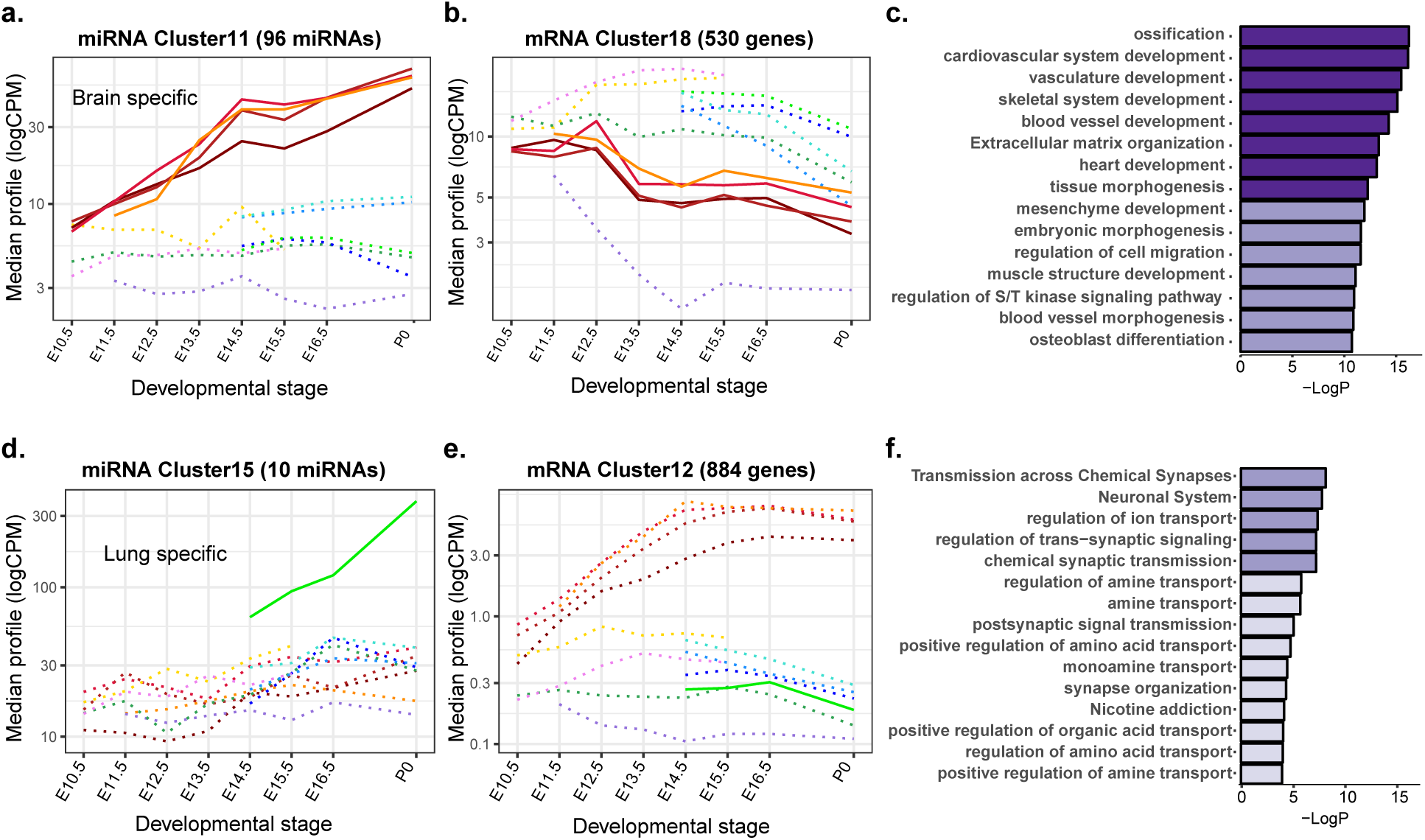
Anti-correlation of microRNA clusters and their developmental gene targets. **(a)** miRNA cluster 11 corresponds to brain-specific miRNAs upregulated during development. **(b)** mRNA cluster 18 genes are highly expressed in other tissues such as limb, cranioface and heart. **(c)** Gene ontology of miRNA cluster 11 targets in mRNA cluster 18 shows enrichments in developmentally important genes with roles outside the brain. **(d)** miRNA cluster 15 increases specifically during lung development. **(e)** mRNA cluster 12 gene expression goes up in brain. **(f)** GO analysis of miRNA cluster 15 targets in mRNA cluster 12 revealed brain specific terms such as synaptic transmission.

## DISCUSSION

In this study we provide a comprehensive resource of miRNA expression dynamics across mouse developmental stages in multiple tissues. Our catalogue of tissue and developmental stage specific miRNAs provides a valuable resource for elucidating the role of miRNAs and highlighting certain key properties of miRNAs during mouse development. we detected only ~25% of the annotated miRNAs in mouse (~60% of miRNAs annotated as highly confident) in the 16 different tissues that are representative of major organ systems during mouse embryogenesis. This result suggests that only a subset of miRNAs might be involved in regulating gene expression during mouse development with the remaining either expressed in other tissues or more likely expressed later in post-natal development and adult tissues (Ludwig et al., 2016). There is also little variability in the number of miRNAs detected per tissues with the heart and nervous system tissues exhibiting the highest number of detected miRNAs. Interestingly, the miRNA output of most embryonic samples is dominated by the expression of a few highly expressed miRNAs that usually consist of non-tissue-specific and ubiquitously but highly expressed miRNAs, which matches reports from human and mouse cell types (De Rie et al., 2017).

Although tissue specificity of miRNAs has been well studied and well reported in multiple model organisms (Gao et al., 2011; Lagos-Quintana et al., 2002; Ludwig et al., 2016), a comprehensive study of the dynamics of such tissue-specific miRNAs across mouse development was lacking. Our analysis fills this knowledge gap. We show that most of the tissue-specific miRNAs are dynamically regulated across development, with different subsets of miRNAs in the brain and heart expressed at different levels during embryonic development.

We provide evidence that the tissue-specific expression of a subset of miRNAs is conserved in human and mouse although the overall transcriptional programs are known to have considerably diverged in the two species (Yue et al., 2014). Although the number of one-to-one miRNA orthologs in human and mouse is low as a fraction of the known miRNAs in each species (~20% of annotated miRNAs in human have one-to-one orthologs in mouse), we show that the tissue-specific expression patterns of the miRNA orthologs closely resemble the overall patterns observed in each individual species. The conservation of miRNA expression in human and mouse tissues is driven by core sets of miRNAs. We show that the expression of tissue-specific miRNAs is well conserved in some tissues (brain, muscle, and lung) while less conserved in other tissues (liver). The fraction of conserved miRNAs is significantly lower than the number of conserved genes between human and mouse (Herrero et al., 2016), which suggests that miRNAs are evolving more frequently.

Finally, the clustering of the miRNAs based on the dynamics of their expression in different tissues allowed us for a unique opportunity to study the functionality and role of these miRNAs in a cooperative way. This approach revealed that some of these tissue specific clusters of miRNAs likely act as suppressors of genes involved in the development of other tissues than those in which the cluster of miRNA is expressed. Many of the target genes are transcription factors that are themselves important for mouse development, which strongly suggests that post-transcriptional regulation needs to be incorporated into models of transcriptional regulation being built from ChIP-seq, open chromatin, and mRNA expression data. The availability of microRNA expression levels in matching tissues and time points of the Mouse ENCODE dataset of embryonic development provides a unique opportunity to integrate the analysis of microRNAs with other functional genomic data used to build the Mouse Encyclopedia of DNA Elements.

## SUPPLEMENTARY INFORMATION

### 1. Materials and Methods

#### Experimental Methods

##### microRNA-seq from mouse embryonic tissues

###### Mouse Embryonic Tissue Acquisition

A detailed protocol for tissue acquisition used for this study can be found at:

https://www.encodeproject.org/documents/631aa21c-8e48-467e-8cac-d4Oc875b39l3/@@download/attachment/Tissue_Excision_Protocols_112414.pdf

###### RNA Isolation

Total RNA was obtained using mirVana miRNA isolation kit and protocol:

https://www.encodeproject.org/documents/f0cc5a7f-96a5-4970-9f46-3l7cc8e2d6a4/@@download/attachment/cms_O55423.pdf

The protocol for genomic DNA removal:

https://www.encodeproject.org/documents/428al84d-7fal-4599-9d8d-749c2eba7edd/@@download/attachment/cms_O5574O.pdf

###### Library Construction

The construction of microRNA-seq libraries was based on the previously published protocol (Roberts et al, 2015, Nucleic Acid Research) with some minor modifications listed below and without the highly abundant miRNA blocking step. Briefly, 500ng of total RNA with RIN (RNA integration number) higher than 9.0 was used as input material, together with spike-in control. 3’adapter was ligated to the sample with T4 RNA ligase 2, truncated (NEB), then reverse transcription primer was annealed to the 3’adapter in order to reduce the 5’ and 3’ adapter dimer. After that, 5’ adapter was ligated to the product with T4 RNA ligase 1 (NEB). Here, we used a pool of four multiplex 5’ adapters. At the end of the 5’ adapter, there is a six-nucleotide spacer, which was present as the first six nucleotides in read 1 of the sequence data in order to provide base diversity during the crucial first cycles. Ligation product was reverse transcribed with Superscript II (Invitrogen) and the cDNA was further amplified using Phusion high-fidelity PCR master mix (NEB). Primers used at the PCR stage introduce a barcode, used later for sample demultiplexing. PCR products were purified with Ampure XP beads (Beckman Coulter). To get rid of adapter-dimer and the other non-miRNA product, size selection of the microRNA-seq libraries was performed using 10% TBE-urea polyacrylamide gel (Bio-Rad) in hot (70C) TBE running buffer for 45 mins. The 140-nt denatured microRNA-seq library band was excised, eluted from the gel slice, precipitated by isopropanol and resuspended with 10ul EB buffer (QIAGEN). Library concentration was determined with Library Quantification Kit (KAPA Biosystems). The DNA Bioanalyzer assay is unable to show the accurate profile of the library and was not employed.

###### Sequencing

The microRNA-seq libraries were sequenced as 50 bp single-end reads on an Alumina HiSeq2OOO sequencer.

##### short RNA-seq from human fetal tissues

Detailed protocol for short RNA-seq library construction can be found at https://www.encodeproject.org/ by searching each sample accession ID.

##### NanoString from mouse embryonic tissues

The samples were prepared with NanoString human miRNA kit version 2.1 (based on miRBase v.18) following its protocol. In short, 100ng total RNA was used as starting material. Together with “spike-in” positive and negative controls, each target miRNA was ligated to a specific miRNAtag molecule and the chimeric miRNA:miRNAtag molecule was hybridized with fluorescent-labeled probes overnight. The miRNA:miRNAtag chimeric molecule is long enough to ensure the efficiency and specificity of probe hybridization. After the samples were processed in NanoString nCounter PrepStation to remove unhybridized probes, they were immobilized and aligned in scanning cartridges and scanned in NanoString nCounter digital analyzer with the maximal resolution setting to achieve the counts of each individual target miRNA molecule recognized by probe.

#### Data processing and analysis

##### Adapter trimming of mouse microRNA-seq and human short RNA-seq reads

Due to small size of miRNAs (<30 nt), adapter trimming of raw sequencing reads was an important step before mapping.

###### Mouse microRNA-seq read adapter trimming

We used Cutadapt v.1.7.1 with Python 2.7 to sequentially trim 5’ and 3’ adapters from raw reads. The 3’ and 5’ (a mixture of 4 sequences) adapter sequences are as follows:

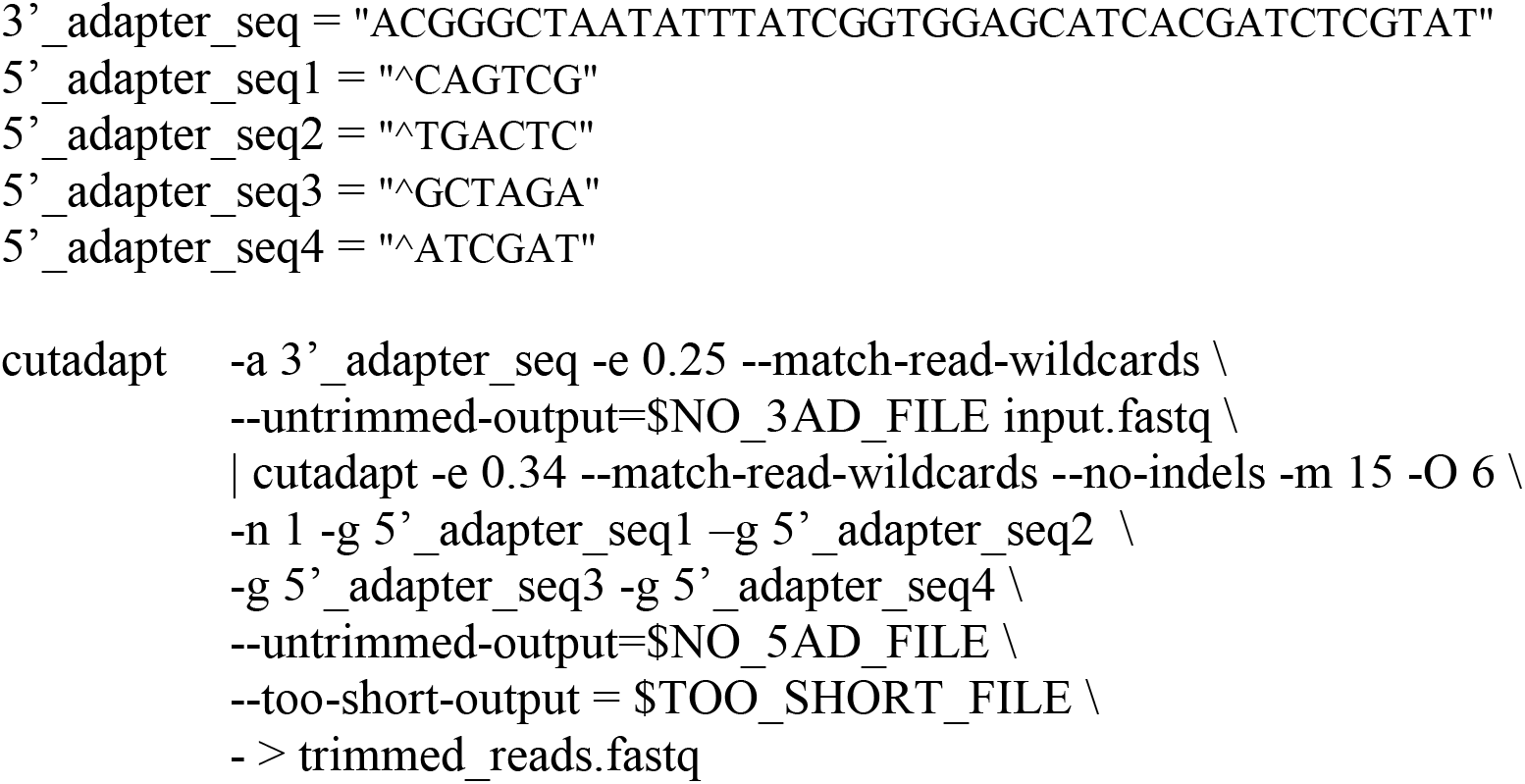

###### Human short RNA-seq read adapter trimming

Reads were initially trimmed for TGGAATTCTC adapters and Ns with cutadapt with parameters: -m 16 --trim-n. In the case of polyA adapters, three additional parameters were used: -a A{10} -e 0.1 -n 10 (to iteratively remove longer polyA tails).

##### Mapping of mouse microRNA-seq reads

Trimmed reads were mapped to the mouse miRBase v.22 mature miRNA sequences with STAR v2.4.2a with parameters:

--runThreadN 16 --alignEndsType EndToEnd --outFilterMismatchNmax 1 --

outFilterMultimapScoreRange 0 --quantMode GeneCounts --outReadsUnmapped Fastx -- outSAMtype BAM SortedByCoordinate --outFilterMultimapNmax 10 --outSAMunmapped Within

--outFilterScoreMinOverLread 0 --outFilterMatchNminOverLread 0 --outFilterMatchNmin 16 -- alignSJDBoverhangMin 1000 --alignIntronMax 1

##### Mapping of human short RNA-seq reads

Trimmed reads were mapped to the human genome (assembly hg38) with STAR v2.5.1b with parameters: --outFilterMultimapNmax 10 -- outFilterMultimapScoreRange 0 --outFilterScoreMinOverLread 0 -- outFilterMatchNminOverLread 0 --outFilterMatchNmin 16 --outFilterMismatchNmax 1 -- alignSJDBoverhangMin 1000 --alignIntronMax 1.

Finally, miRNA hairpins from GENCODE v25 were quantified by summing the reads that have 100% overlap with the hairpins.

##### Processing of mouse NanoString data

NanoString raw data for all samples were processed with NanoStringNorm v.1.2.1 (Waggott, 2012) using the function “NanoStringNorm” with the following parameters: CodeCount = ‘geo.mean’, Background = ‘max’, SampleContent = ‘top.geo.mean’, round.values = TRUE, take.log = FALSE.

##### Differential analysis of human short RNA-seq data from human

miRNA hairpins specific of a given tissue in human or in mouse were identified with the glmQLFit() and glmQLFTest() function() from the R package edgeR. We used a deviation coding system for contrasts which compares one tissue against all the others, with the function constr.sum(). P-values were adjusted for multiple testing with the Benjamini-Hochberg correction. Only hairpins with FDR<0.01 and a fold-change > 2 were considered tissue-specific.

##### Identification of orthologous miRNAs in mouse and human

Orthologous miRNAs between human and mouse were identified through genomic alignment. Human miRNAs were lifted over to the mouse genome and vice versa, with a minimum overlap requirement of 50%. If a human miRNA maps within 10kb of a mouse miRNA, and that mouse miRNA also maps within 10kb of the initial human miRNA, those miRNA are defined to be in reciprocal orthologous relationship.

##### Generation of ab initio transcripts models from mRNA-seq reads

mRNA-seq reads were mapped to the mouse genome (assembly mm10) using STAR v.2.4.2a with the following parameters:

--genomeDir star.gencodeM10.index --readFilesIn $fastqs –sjdbGTFfile gencode.vM10.annotation.gtf --readFilesCommand zcat --runThreadN 8 -- outFilterMultimapNmax 20 --alignSJoverhangMin 8 --alignSJDBoverhangMin 1 -- outFilterMismatchNmax 999 --outFilterMismatchNoverReadLmax 0.04 --alignIntronMin 20 -- alignIntronMax 1000000 --alignMatesGapMax 1000000 --outSAMunmapped Within -- outFilterType BySJout --outSAMattributes NH HI AS NM MD XS --outSAMstrandField intronMotif --outSAMtype BAM SortedByCoordinate --sjdbScore 1

The alignments to the genome were assembled into ab initio transcripts using StringTie v.1.2.4 using the following parameters:

-G gencode.vM10.annotation.gtf -c 3 -p 8

The transcript models for each sample were merged into a single GTF file using StringTie with the following options:

stringtie --merge -G gencode.vM10.annotation.gtf -o merged.gtf

-m 200 -F 1.0 -p 8

Single exon transcripts with no strand information were excluded. The expression levels of the GENCODE M10 and the new StringTie model transcripts were obtained using RSEM v1.2.25 with the following parameters:

rsem-calculate-expression --star --star-path ~/STAR-STAR_2.4.2a/bin/Linux_x86_64/ -p 10 -- gzipped-read-file fastqs RSEM_Index_GENCODE_M10_Plus_StringTieModels

##### Prediction of novel miRNAs from mouse microRNA-seq and human short RNA-seq reads

###### Mouse

All the trimmed microRNA-seq reads of samples for the same tissue were pooled and novel miRNAs predicted using mirdeep2 v2.0.0.8. The parameters used were:

mapper.pl trimmed_reads.fastq -e –p mm10

miRDeep2.pl processed.reads.fastq mm10.fasta mapped.arf

We used miRBase v.21 mature and hairpin annotations for mouse and ENSEMBL v.85 rat hairpin annotations for the miRDeep2.pl step above. Novel miRNAs with a score of 4 or higher (corresponding to 70% or higher true positive confidence level), independently identified in at least two tissues, not overlapping GENCODE M10 annotated miRNAs and the genomic repeat regions, and expressed at 2 CPM minimum in at least one sample (both replicates) were kept for downstream analysis.

##### Time-series analysis of mouse microRNA-seq and NanoString data

Time series analysis of the mouse microRNA-seq time-course was performed using maSigPro_1.48.0 in R 3.4.4. Briefly, each tissue (12 in total) that were assayed in at least two developmental time points were analyzed using a degree 3 and maSigPro functions “p.vector(data, design = design.matrix, counts = TRUE)”, “T.fit(p.vector_output, alfa = 0.01)”, and “get.siggenes(T.fir_output, rsq=0.7, vars=“all”)”. Different number of clusters (k) were tested to find the best meaningful number of clusters by contrasting the newly formed clusters at each step of k with the previous ones using the command “see.genes(get$sig.genes, k = …)”. The best results for miRNAs were given with k=16. The median profiles of the genes (standard errors denoted for the replicates) were plotted using ggplot2 package. The code used to generate these clusters and plots can be found at:

<https://github.com/sorenar/mouse_embryonic_miRNAs/blob/master/miRNA_maSigPro.R> Time-series analysis of mouse mRNA-seq data:

Time series analysis of the mouse mRNA-seq time-course was performed similar to the clustering of miRNAs. In this case higher number of clusters (k = 20-35) was tested and k=30 was selected as the number that gave the best results. Similarly, the median profiles of these clusters were plotted using ggplot2. The code used to generate these clusters and plots can be found at:

<https://github.com/sorenar/mouse_embryonic_miRNAs/blob/master/mRNA_maSigPro.R> Tissue specificity analysis of individual miRNAs:

The miRNAs tissue specificity was determined using a tissue specificity index as described by Ludwig 2016:

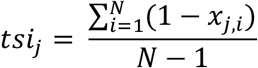

In order to prevent any biases introduced by multiple tissues of neural origin we excluded the samples from hindbrain, midbrain and neural tube and used only the forebrain samples for tissue specificity study of individual miRNAs. Also since some of the tissues have missing data from the initial stages (due to the fact that they start development later), we decided to restrict the number of tissues considered for TSI calculations to cranioface, forebrain, heart, limb and liver for the first 4 stages and to the cranioface, forebrain, heart, limb, liver, stomach, cranioface and limb for the last 4 stages.

##### Identification of tissue specificity of the miRNA clusters

The average expression of miRNAs in each cluster was calculated for each sampling point (for each tissue at each of the time points). Then the standard deviation of each cluster was calculated for each tissue across different time points. The standard deviations obtained for different tissues were then scaled for each miRNA cluster and the tissues with positive values were considered as the tissue specificity of the miRNA cluster. The code to create this tissue specificity matrix and the corresponding plot is available at:

<https://github.com/sorenar/mouse_embryonic_miRNAs/blob/master/PartialCorrelation.R> Building the partial correlation matrix:

Similar to the method explained above, the average expression of mRNAs in each cluster were calculated at each tissue at every time point. For each pairs of miRNA-mRNA clusters only the sample points corresponding to the tissues identified as specific to the miRNA cluster were used to find the Pearson correlation. The code to generate this partial correlation matrix is provided at:

<https://github.com/sorenar/mouse_embryonic_miRNAs/blob/master/PartialCorrelation.R> Target enrichment analysis using miRNAtap:

The R package, miRNAtap (1.10.0) was used as an ensemble method to compile the predicted targets for each miRNA in our data. miRNAtap relies on five different sources to come up with a list of predicted targets, these sources are: DIANA (Maragkakis et al., 2011), Miranda (Enright et al., 2003), PicTar (Lall et al., 2006), TargetScan (Friedman et al., 2009), and miRDB (Wong and Wang, 2015). We used the command getPredictedTargets(miRNA, species = ‘mmu’,method = ‘geom’,min_src = 3) to obtain the list of predicted targets for miRNA. The parameter “min_src” indicates that if the miRNA has targets that are present in more than “min_src” value, the reported list would be only limited to those targets, otherwise the method will reduce the “min_src” until it gets a list of targets or no target at all.

We compared this ensemble approach with the approach that uses only targetScan to see if we can recover similar results. Most of the enriched interactions (78%) reported by “min_src = 3” method had 50-90% of their targets recovered by TargetScan. However, when we re-build the enrichment matrix using only TargetScan, there are 27 significant interactions, only 3 of which overlapped with the “min-src = 3” method. Furthermore, when we looked at the profile comparison of these significantly enriched interactions, we find that most of the miRNA clusters has lower tissue specificity.

The code for the target enrichment analysis can be found in:

<https://github.com/sorenar/mouse_embryonic_miRNAs/blob/master/TargetEnrichment.R>

##### Gene ontology analysis of significant interactions

For each significant interaction with a negative partial correlation, the list of the target genes in the interaction was compiled. The gene ontology analysis of each of these target list was performed via Metascape (Tripathi et al. 2015) and the plots were generated using the code in:

<https://github.com/sorenar/mouse_embryonic_miRNAs/blob/master/SigIntAnalysis.R>

---

**Supplementary Figure 1:**
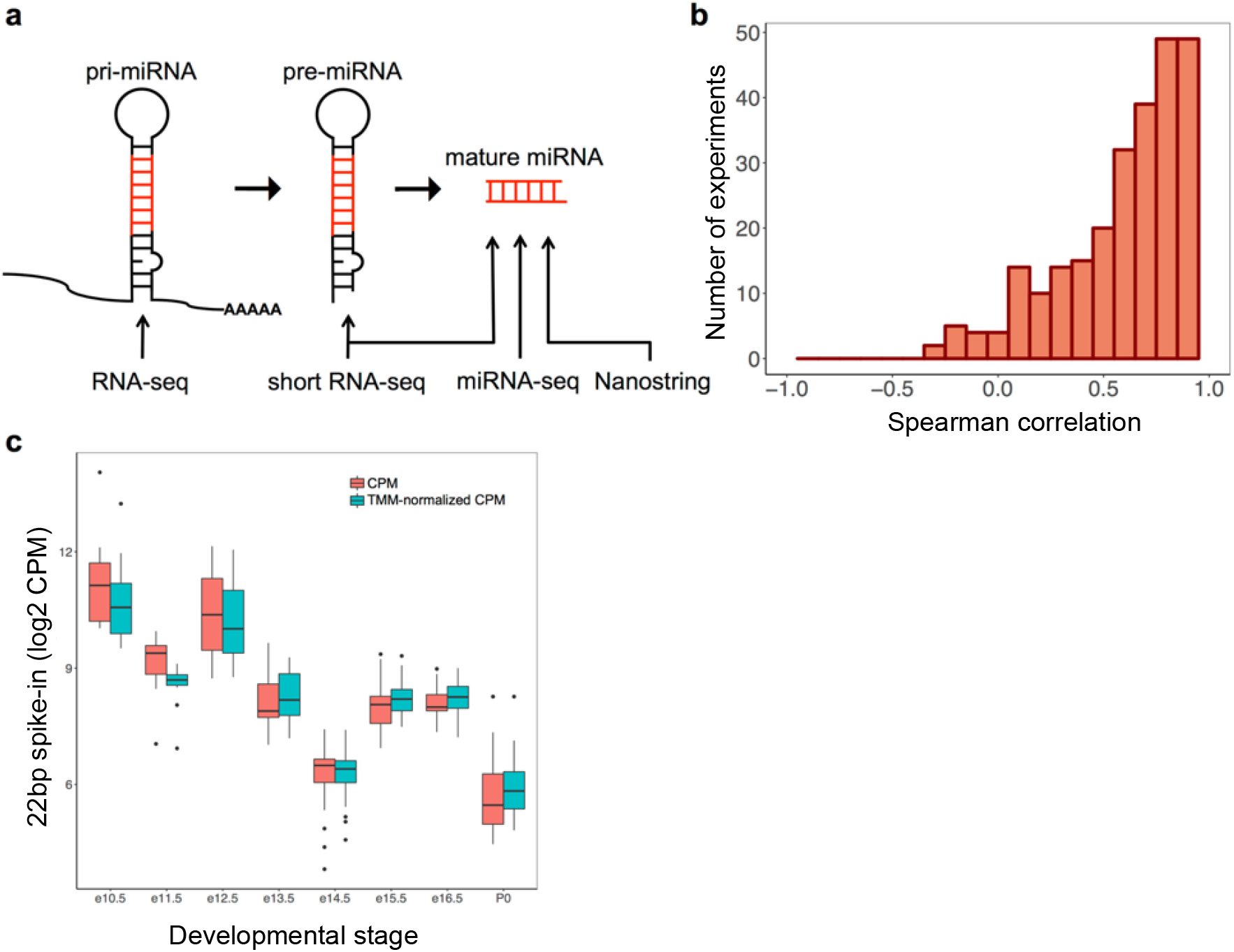
Overview of mouse ENCODE miRNA data sets. **(a)** Primary miRNAs were profiled using mRNA-seq (> 200 nt) in human and mouse, pre-miRNAs and mature miRNAs were profiled in human using short RNA-seq (< 200 nt), and mature miRNAs in mouse were profiled using microRNA-seq (< 30 nt) and NanoString. **(b)** Distribution of Spearman correlations (median = 0.68) between microRNA-seq and NanoString for miRNAs included in the NanoString coreset. **(c)** Expression levels of the 22 bp spike-in in mouse microRNA-seq samples across different stages of embryonic development for non-normalized counts-per-million (CPM, red boxplots) and TMM-normalized CPMs (cyan boxplots).

**Supplementary Figure 2:**
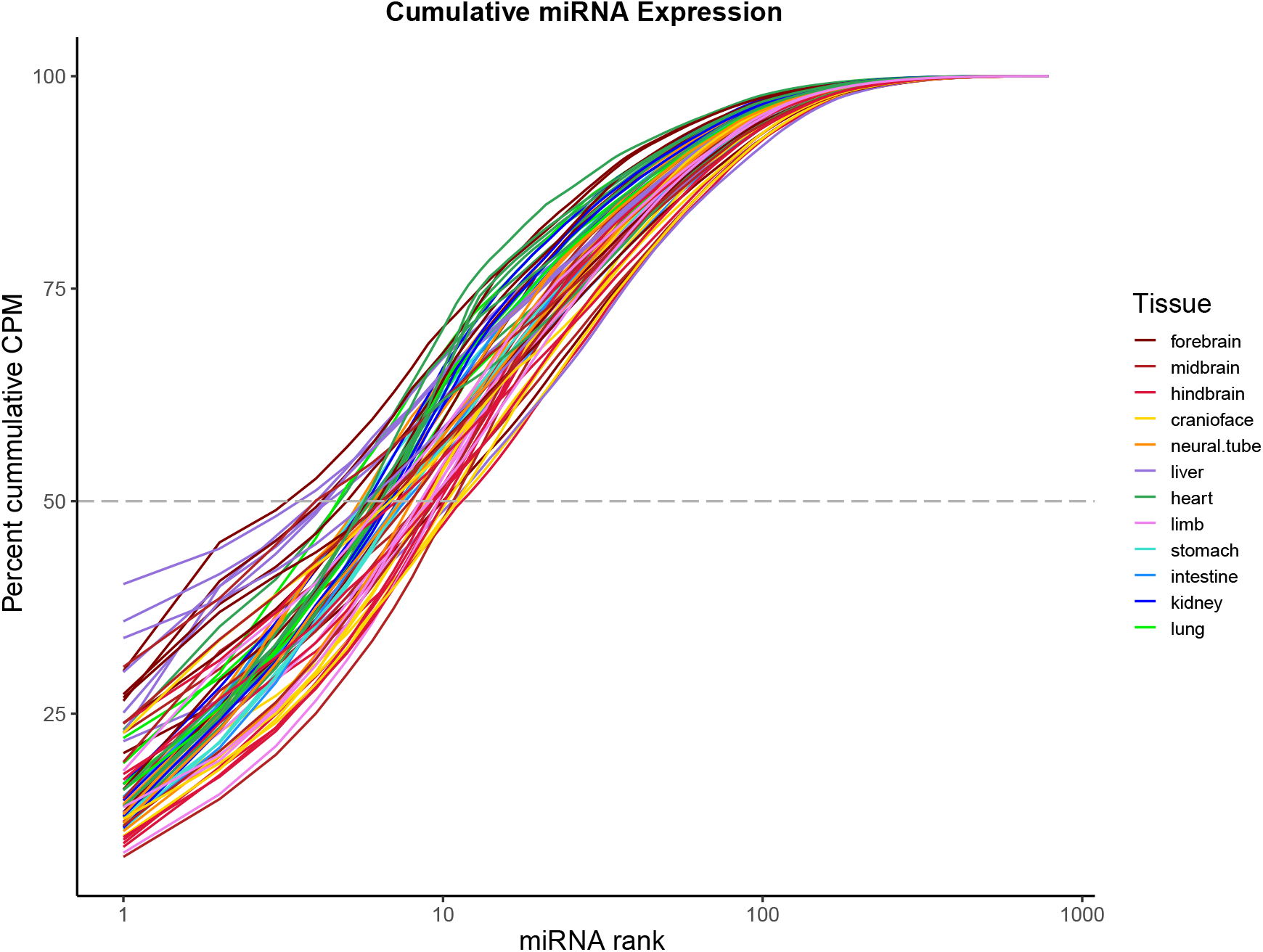
A few highly expressed miRNAs dominate miRNA-seq datasets. Cumulative distribution of sequencing reads, using miRNAs ranked by expression levels. The top 10 miRNAs account for more than half of the miRNA sequencing reads.

**Supplementary Figure 3:**
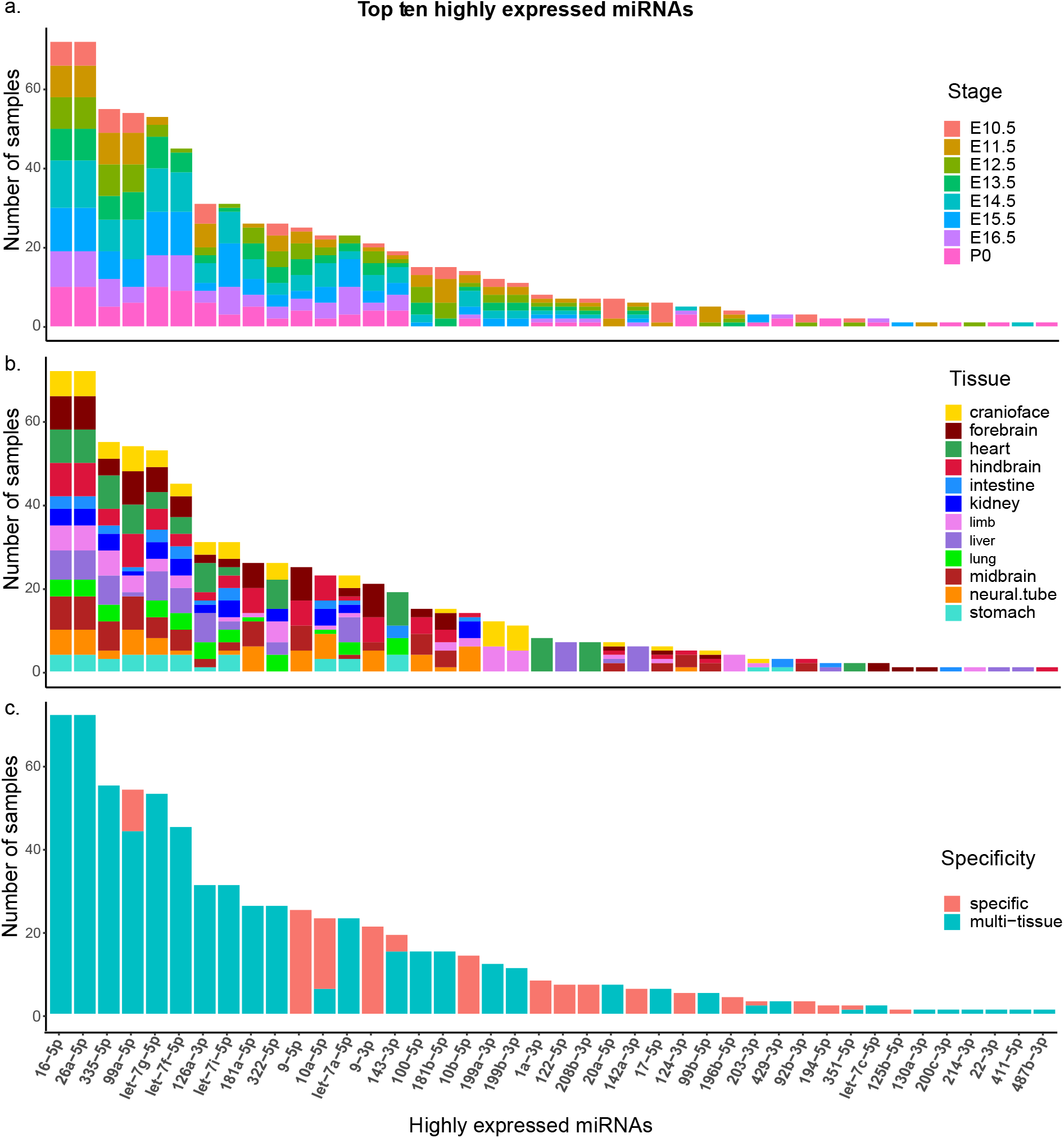
Characterization of the specificity of highly expressed miRNAs. Histograms of the 43 miRNAs that form the top ten highly expressed miRNAs of all the samples, ranked by the number of samples they are highly expressed in; colored by their tissue **(a),** stage **(b)** and specificity **(c).**

**Supplementary Figure 4:**
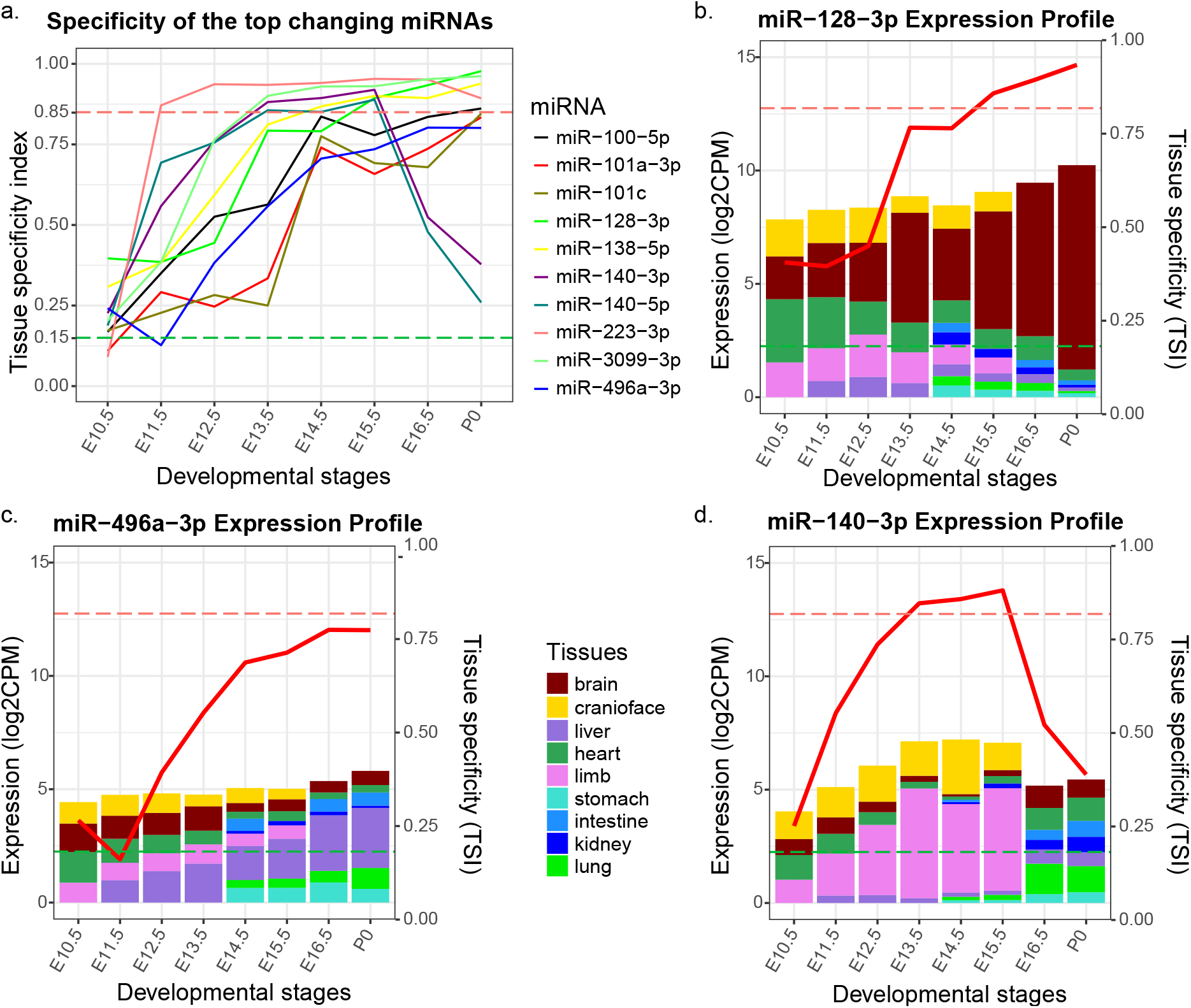
Analysis of miRNAs that change their specificity. **(a)** The tissue specificity profile of ten miRNA with the most change in their specificity through the embryonic development. **(b-d)** The bar graphs show the expression profile of three of these miRNAs: **(b)** miR-128-3p, **(c)** miR-496a-3p, **(d)** miR-140-3p. The red curve traces the tissue specificity (TSI) on the second axis and each bar is colored proportional to the expression of the miRNA in different tissues. The fall in tissue specificity of miR-143-3p at the last two time point may be due to the lack of limb samples for those two time-point.

**Supplementary Figure 5:**
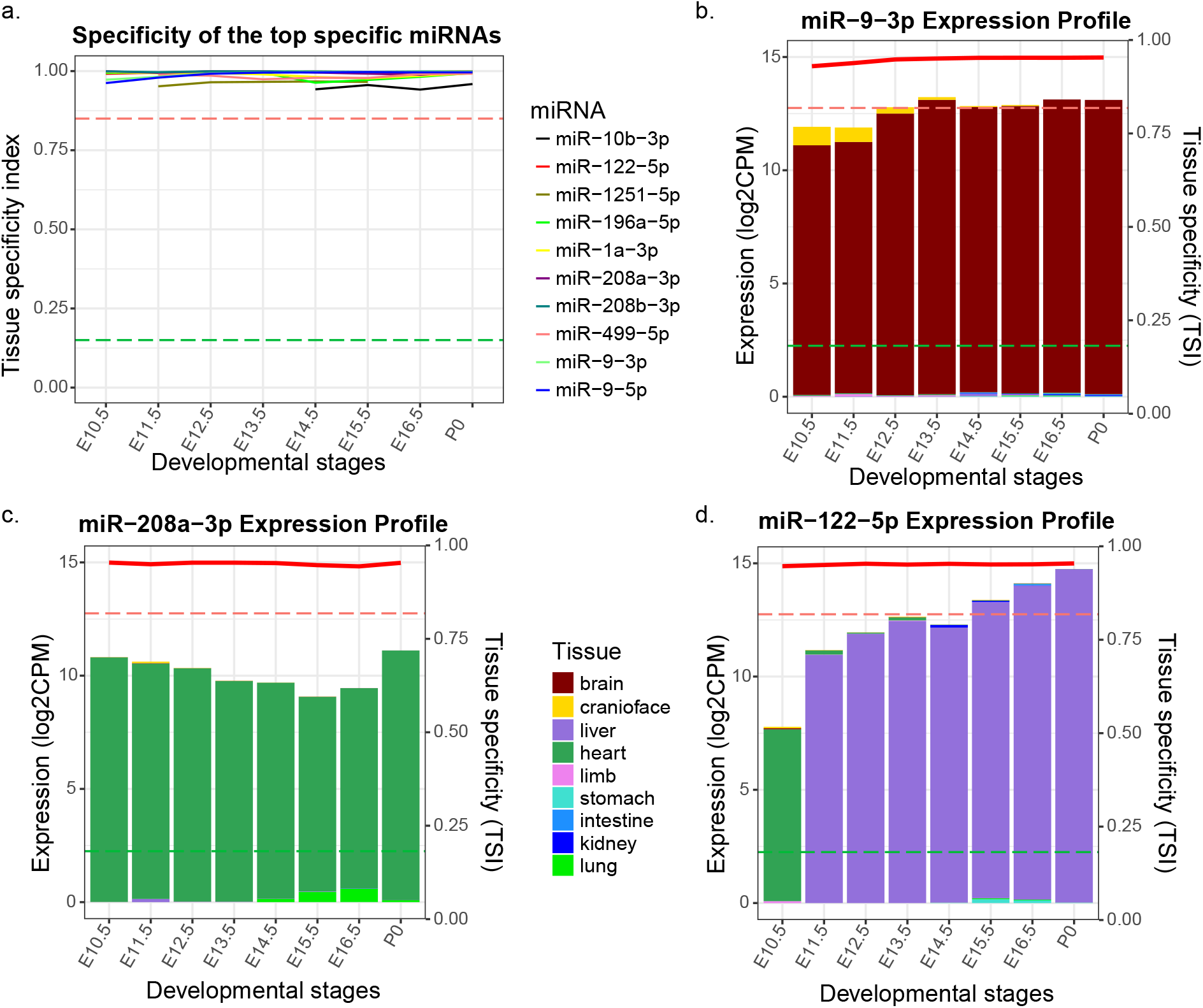
Analysis of miRNAs that are highly specific. **(a)** The tissue specificity profile of ten miRNA with the highest specificity through the embryonic development. **(b-d)** The bar graphs show the expression profile of three of these miRNAs: **(b)** miR-9-3p, **(c)** miR-2O8a-3p, **(d)** miR-122-5p. The red curve traces the tissue specificity (TSI) on the second axis and each bar is colored proportional to the expression of the miRNA in different tissues. Notice that miR-122-5p is highly specific at all stages but is shown as specific to heart at the E10.5 and liver specific at every other stage, possibly due to no liver sample in our data for stage E10.5.

**Supplementary Figure 6:**
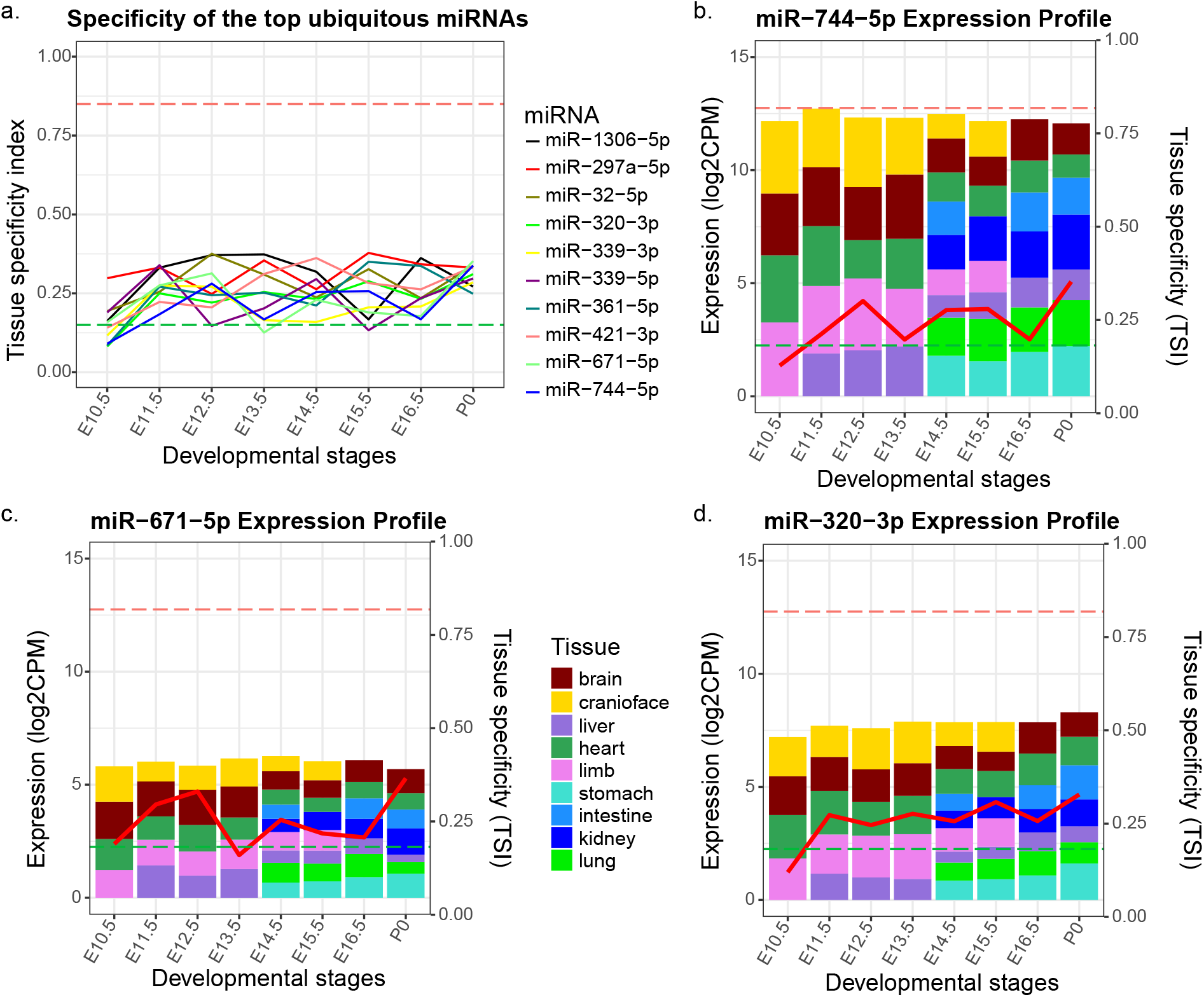
Analysis of miRNAs that are mostly ubiquitous. **(a)** The tissue specificity profile of ten miRNA with the lowest specificity through the embryonic development, **(b-d)** The bar graphs show the expression profile of three of these miRNAs: **(b)** miR-744-5p, **(c)** miR-67l-5p, **(d)** miR-320-3p. The red curve traces the tissue specificity (TSI) on the second axis and each bar is colored proportional to the expression of the miRNA in different tissues.

**Supplementary Figure 7:**
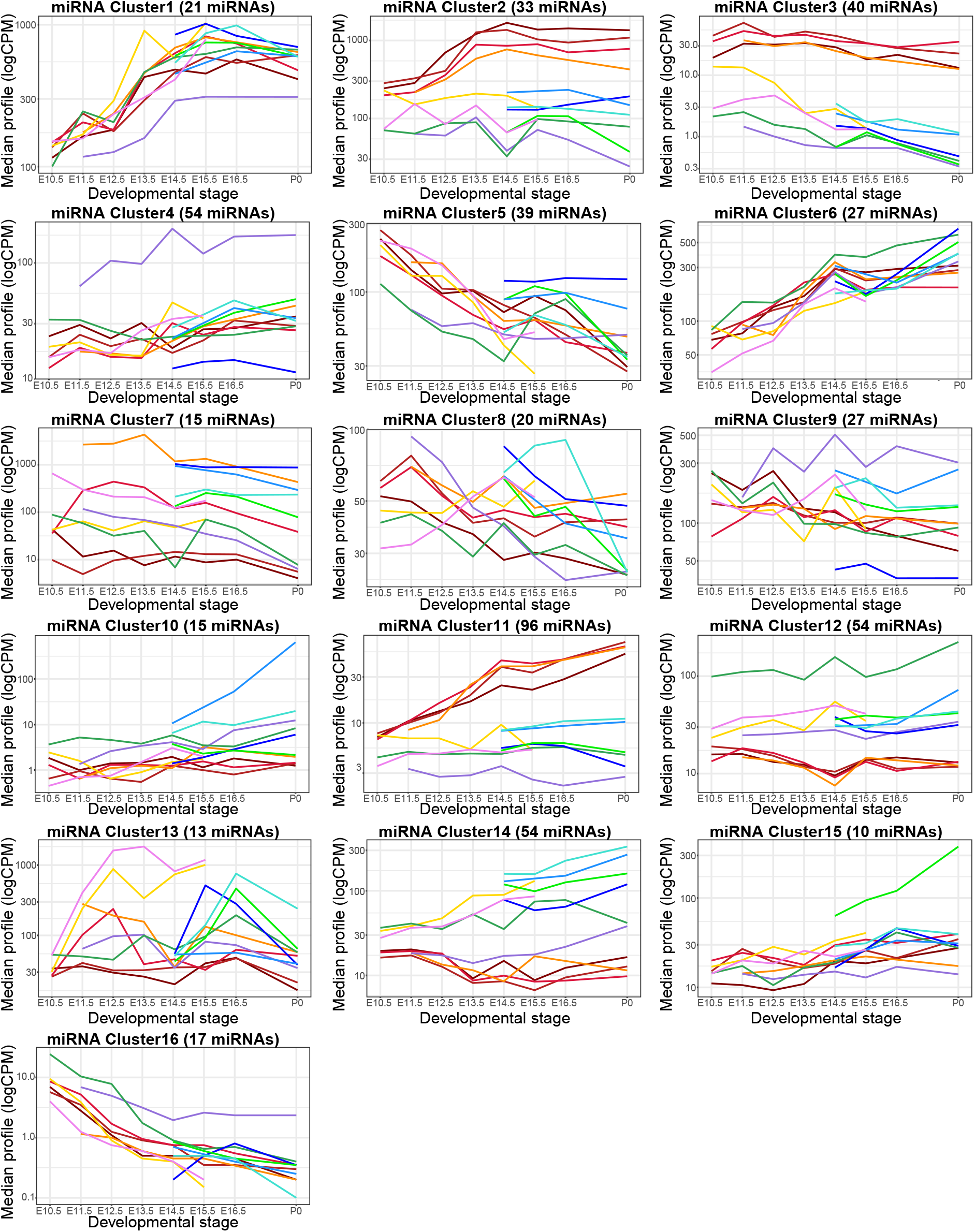
Expression profiles of microRNA-seq miRNA clusters. Median expression levels of 16 clusters of differentially expressed mouse miRNAs that were assayed using microRNA-seq

**Supplementary Figure 8:**
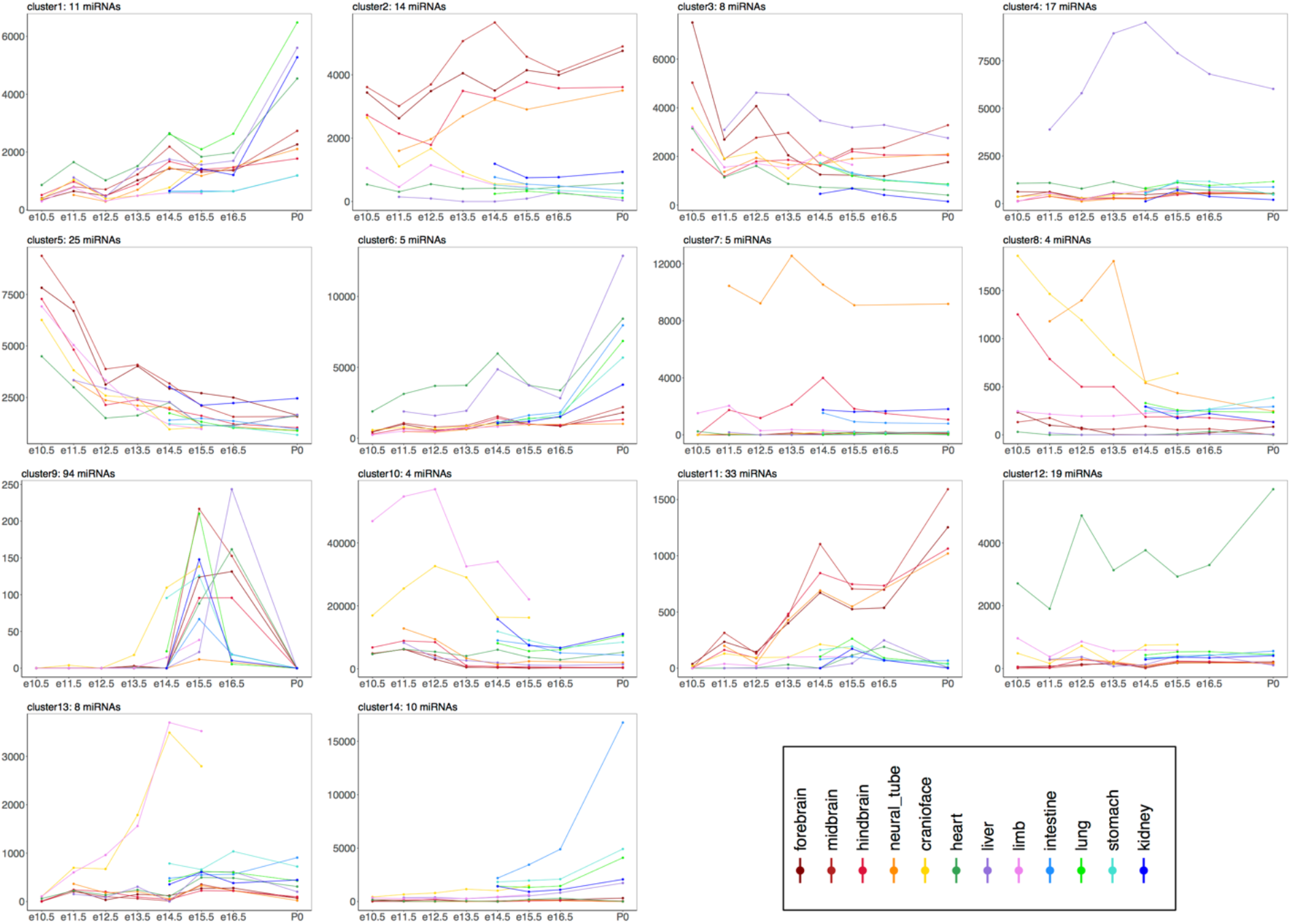
Expression profiles of NanoString miRNA clusters. The median expression profiles of 14 clusters of miRNAs, measured using NanoString in matching samples, that were identified as differentially expressed by the linear regression based algorithm maSigPro.

**Supplementary Figure 9:**
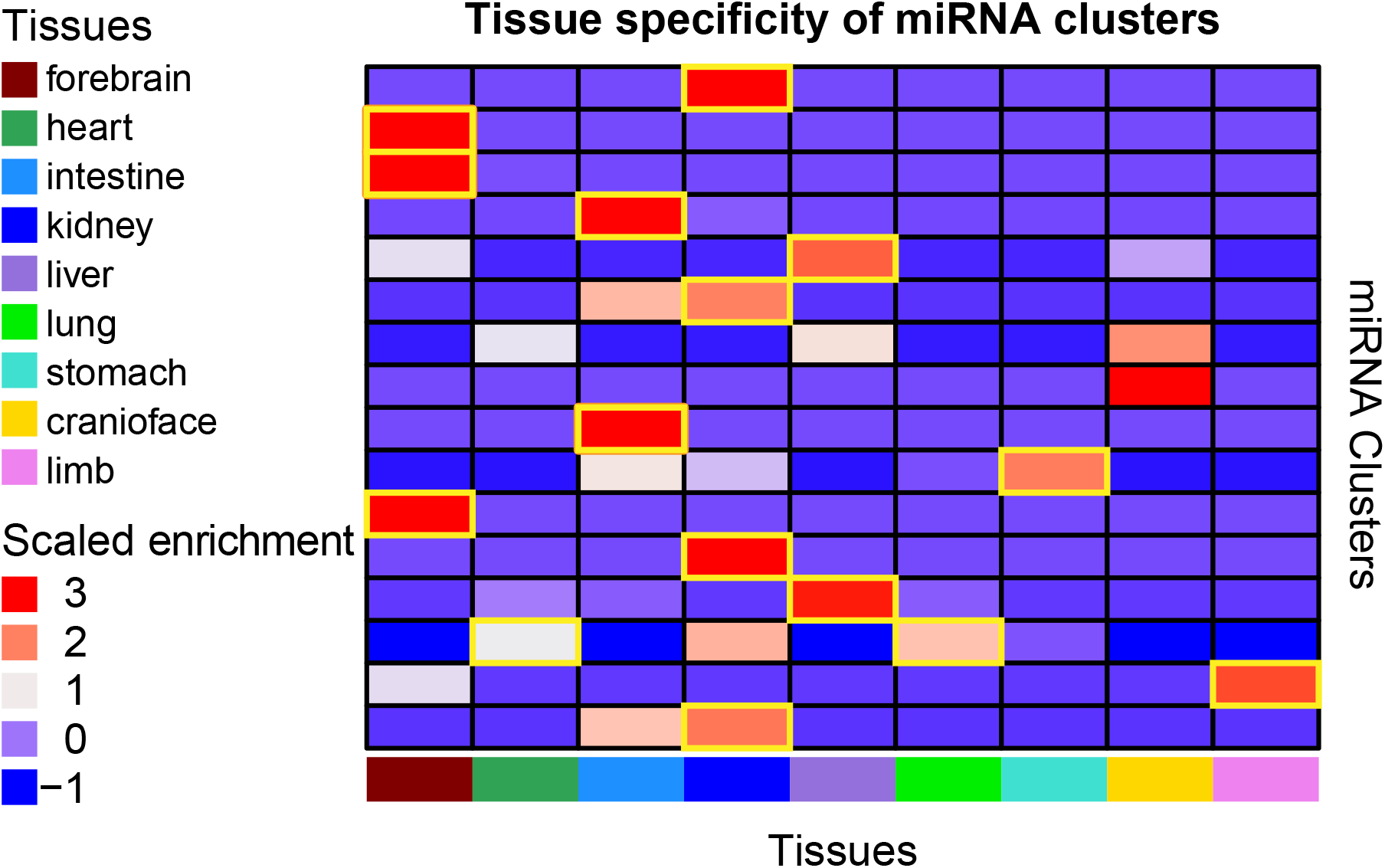
miRNA cluster tissue-specificity enrichment. We further determined the tissue-specificity of the miRNA clusters by enrichment analysis of the tissue-specific miRNAs in each cluster. Tissue-specificity of the majority of the miRNA clusters corresponds with what has been determined using the variance of the average miRNA expression of each cluster within each tissue, with gold boxed clusters indicating where the tissue specificities are concordant with cluster tissue specificity in Fig. 3)

**Supplementary Figure 10:**
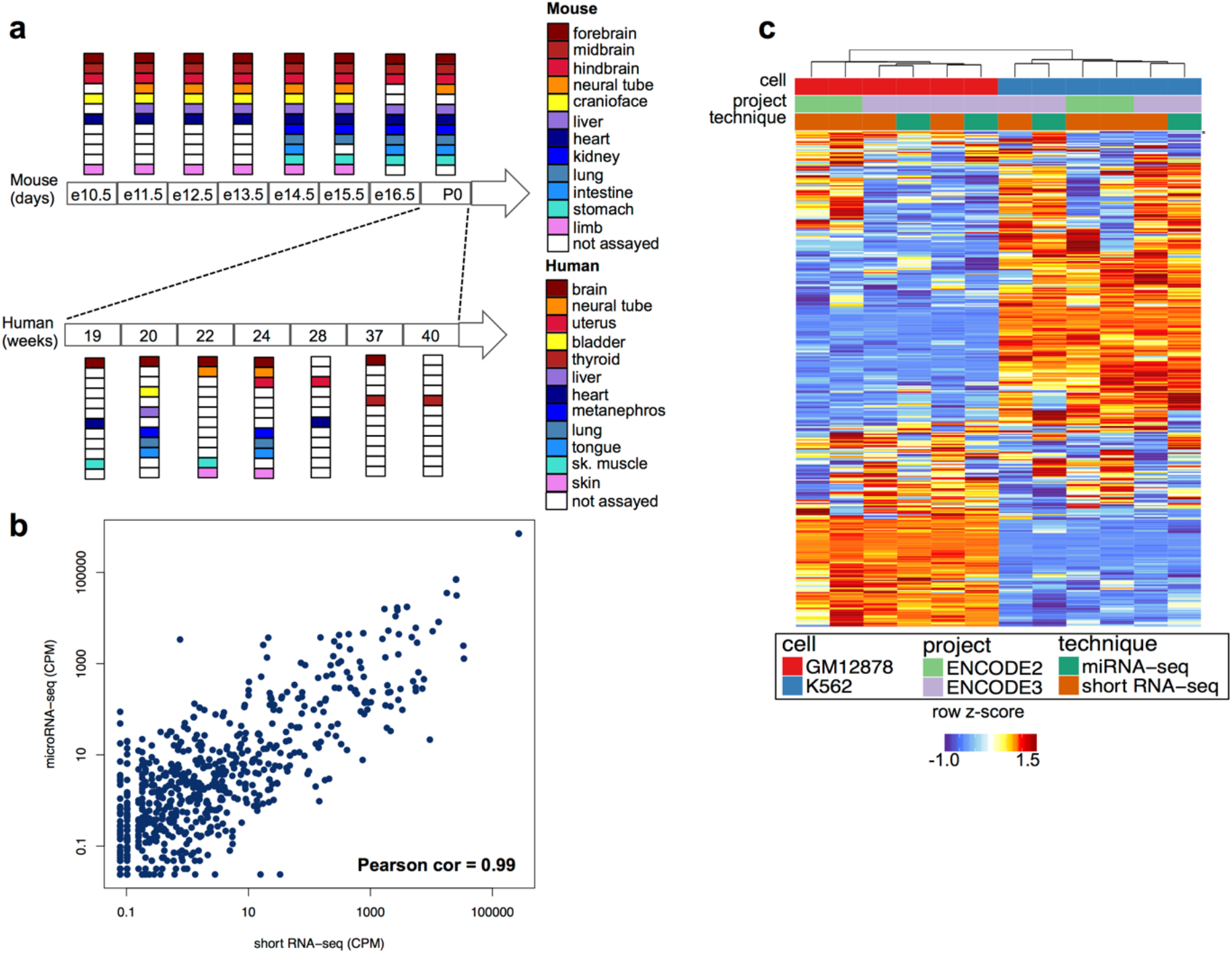
Overview of human ENCODE miRNA data sets. **(a)** Primary tissues representative of major organ systems were profiled in human along various stages of fetal development (weeks 19-40). **(b)** Comparison of normalized miRNA counts for GM12878, profiled using microRNA-seq and short RNA-seq, demonstrates high correlation between the two assays, (c) Heatmap of miRNA normalized counts in GM12878 and K562 cell lines shows that the samples cluster by cell type irrespective of profiling technique used or the date of sample preparation.

**Supplementary Figure 11:**
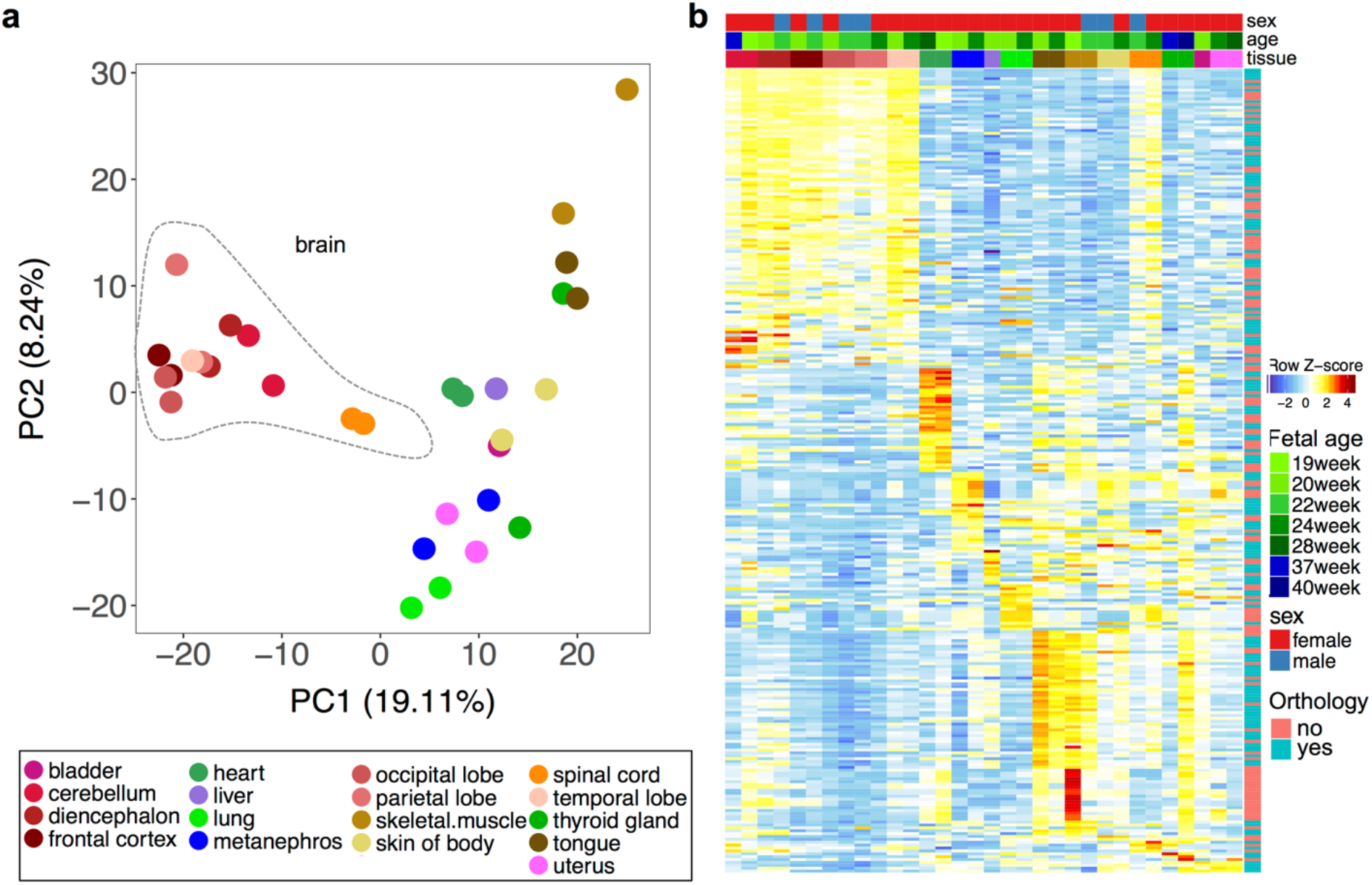
Human fetal development miRNA transcriptome. **(a)** Principal component analysis (PCA) of human tissues reveals distinct miRNA expression patterns in brain samples compared to other tissues. Colors denote different tissues. **(b)** Normalized expression levels of human tissue-specific miRNAs. Differential expression analysis reveals the largest set of differentially expressed miRNAs in brain and muscle samples.

**Supplementary Figure 12:**
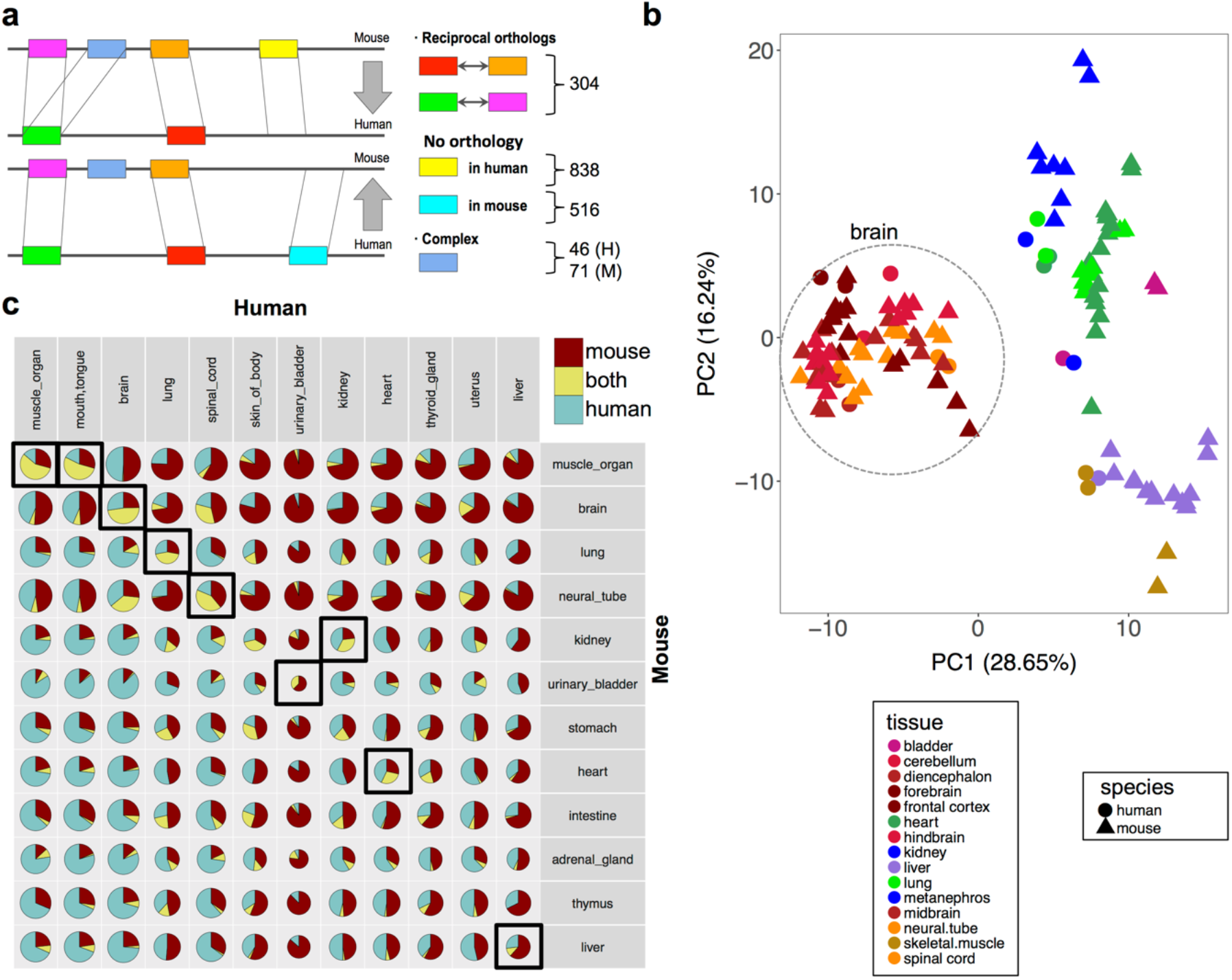
Comparative dynamics of miRNAs during human and mouse development. **(a)** 304 one-to-one orthologous human and mouse miRNAs were identified using reciprocal search. **(b)** Combined principal component analysis (PCA) of human and mouse samples. Triangles and circles denote mouse and human tissues respectively. Tissues are denoted by different colors. **(c)** Intersection of human and mouse tissue-specific miRNAs. For each pair of tissues in human and mouse we report the fraction of tissue-specific miRNAs in mouse only (red), human only (blue), or in both (yellow) within the 304 orthologous miRNAs. The sizes of the pie chart are in proportion to the numbers of tissue-specific miRNAs in the corresponding tissues.

**Supplementary Figure 13:**
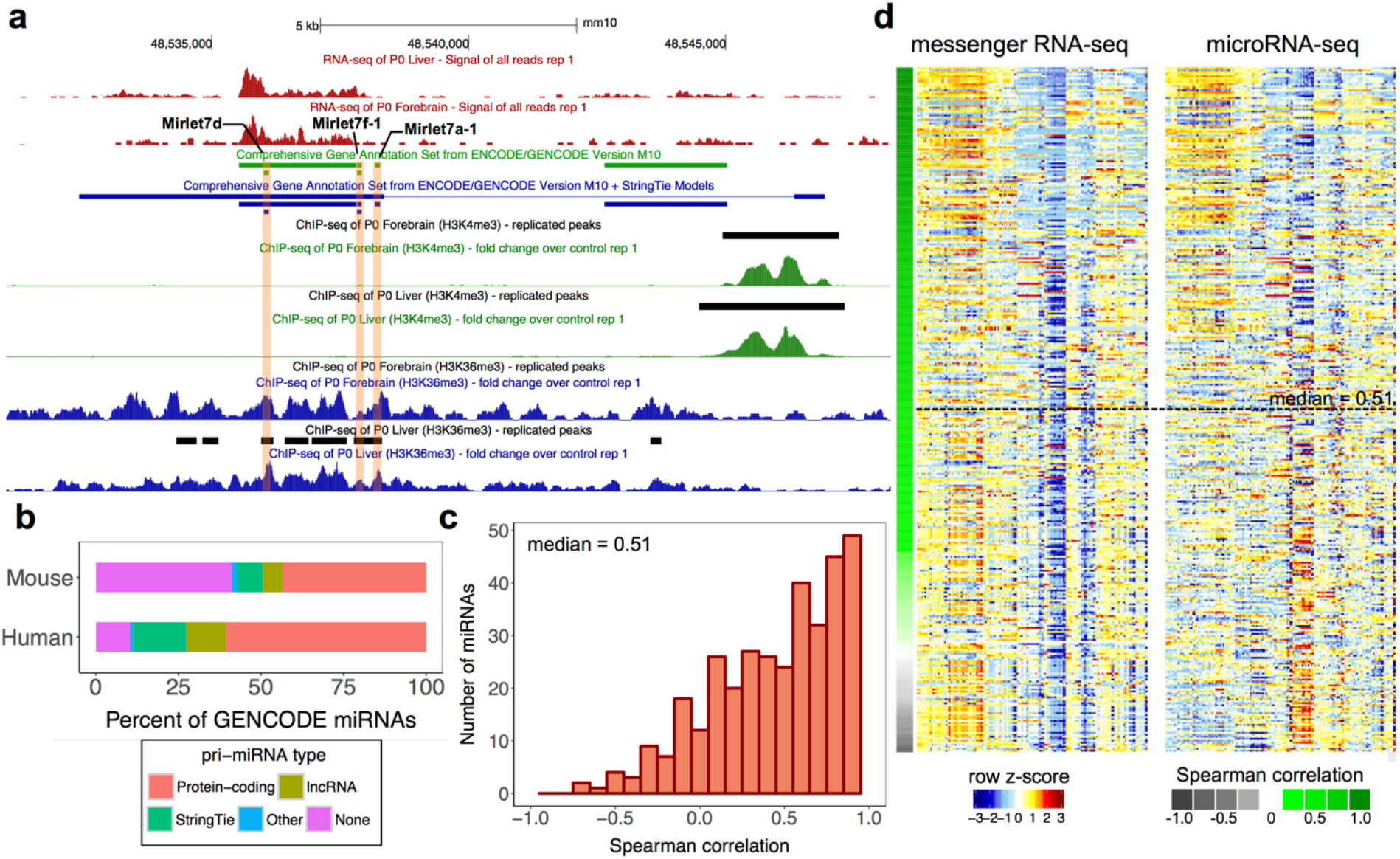
Comparison of miRNAs and their primary transcripts using GENCODE annotations augmented with *ab initio* models. **(a)** A genome browser snapshot of a representative *ab initio* gene model generated using mouse mRNA-seq data for 2 miRNAs (Mir-let7f-1 and Mir-let7a-1) that lack annotated GENCODE v. M10 pri-miRNAs. Additional evidence for the gene models is provided using H3K4me3 and K3K36me3 data in matching mouse samples. **(b)** Distribution of types of pri-miRNAs biotypes (protein-coding, lncRNA, *ab initio* gene models, and others) in mouse and human. Improvements in pri-miRNA annotations using the *ab initio* gene models are denoted by color green in both human and mouse. **(c)** Distribution of Spearman correlation (median = 0.51) among miRNAs and their corresponding pri-miRNAs. **(d)** Comparison of expression levels of pri-miRNAs and their corresponding mature miRNAs. The rows are sorted with decreasing Spearman correlation top to bottom.

**Supplementary Figure 14:**
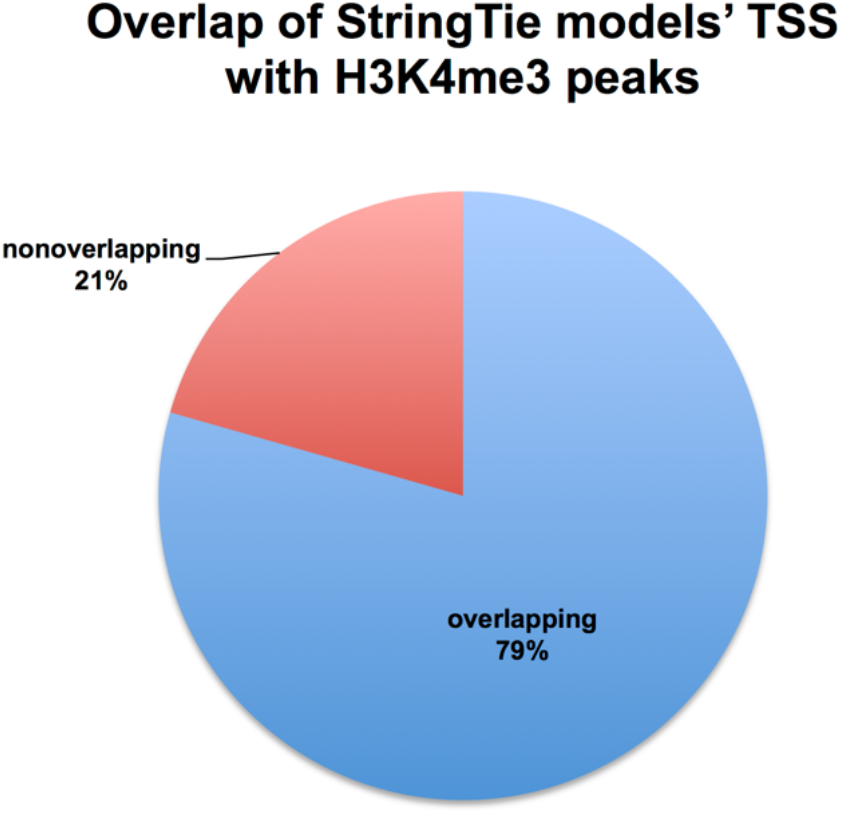
Distribution of StringTie transcript models’s TSS that have overlapping H3K4me3 peaks. Proportion of overlap between the TSS of StringTie *ab initio* transcript models and the H3K4me3 peaks in matching samples. The TSS of the transcript models were defined as ±250 bp regions from the 5’-end of the transcripts.

**Supplementary Figure 15:**
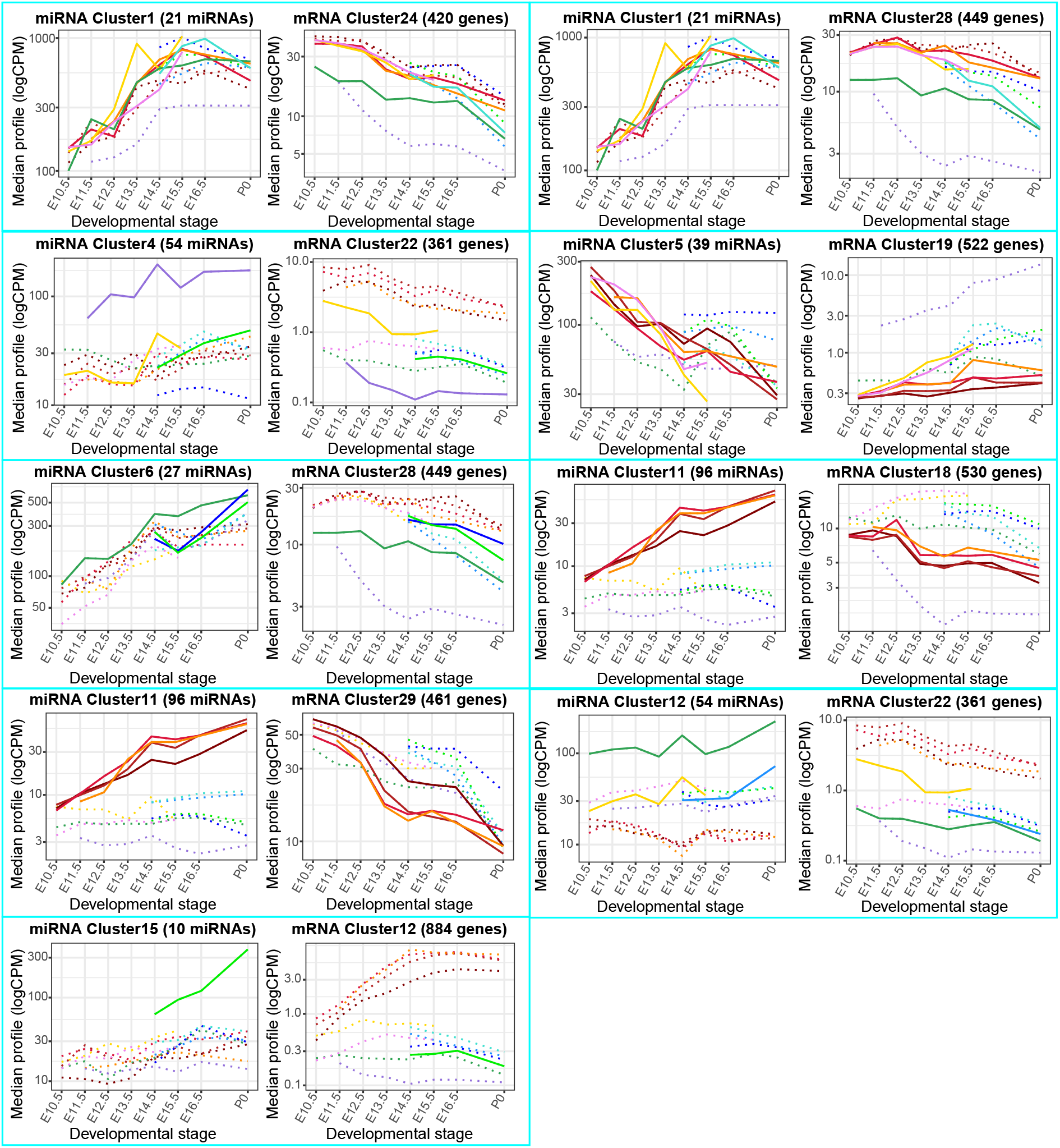
Expression profiles of all the significant interactions. The miRNA and mRNA expressions for the miRNA and mRNA clusters in the identified significant interactions.

